# Bichromatic exon-reporters reveal voltage-gated Ca^2+^-channel splice-isoform diversity across *Drosophila* neurons *in vivo*

**DOI:** 10.1101/2024.11.08.622738

**Authors:** Touhid Feghhi, Roberto X. Hernandez, Olena Mahneva, Carlos D. Oliva, Gregory T. Macleod

## Abstract

Every neuron contains the same genomic information but its complement of proteins is the product of countless neuron-specific steps including pre-mRNA splicing. Despite advances in RNA sequencing techniques, pre-mRNA splicing biases that favor one isoform over another are largely inscrutable in live neurons *in situ*. Here, in *Drosophila*, we developed bichromatic fluorescent reporters to investigate alternate splicing of *cacophony* – a gene that codes the pore-forming α_1_-subunit of the primary neuronal voltage-gated Ca^2+^ channel (VGCC). These reporters reveal a neuron-specific pattern of exon biases, including stereotypical differences between neurons of the same neurotransmitter type and ostensibly the same function. Information about exon splicing biases of individual neurons *in vivo* provides clues to the role of VGCC motifs and the role of those neurons in the context of local circuits. The application of this technology to a large gene such as *cacophony* provides a precedence for effective exon-reporter design for other *Drosophila* genes.

## Introduction

Voltage gated Ca^2+^ channels (VGCCs) are essential conduits for the Ca^2+^ that triggers neurotransmitter release at chemical synapses and are therefore some of the most influential proteins in the nervous system. The identity of a VGCC’s pore-forming α_1_-subunit (Fig. 1A), alternate splicing, post-translational modification, auxiliary subunits (Fig. 1B), and other associated proteins, determine its function and placement. The human genome contains ten genes for α_1_-subunits alone, and a small subset of these genes is expressed in individual neurons (Catterall et al., 2005). *Drosophila* has three homologs of these α_1_-subunit genes, but only one gene, *cacophony*, represents the Ca_v_2 family of α_1_-subunits (Littleton & Ganetzky, 2000). Never-the-less, further diversity in α_1_-subunit form and function can arise through alternate splicing, and *cacophony* has 18 annotated splice isoforms (http://flybase.org/; Fig. 1C). Alternate splicing confers diverse biophysical properties to *cacophony* (Bell et al., 2024), furthermore, cacophony’s roles in synapse assembly (Ghelani & Sigrist, 2018) and synaptic homeostatic plasticity (Frank et al., 2006) are also likely to depend on alternate splicing. Cell specific information about isoform bias can offer insight into the functional role of motifs within the α_1_-subunit, a neuron’s function in the context of a circuit, and risk factors in disease processes (Lipscombe & Andrade, 2015). However, while antibodies, toxins and pharmacological agents have been used to glean isoform specific information, they have many limitations.

**Figure 1:**
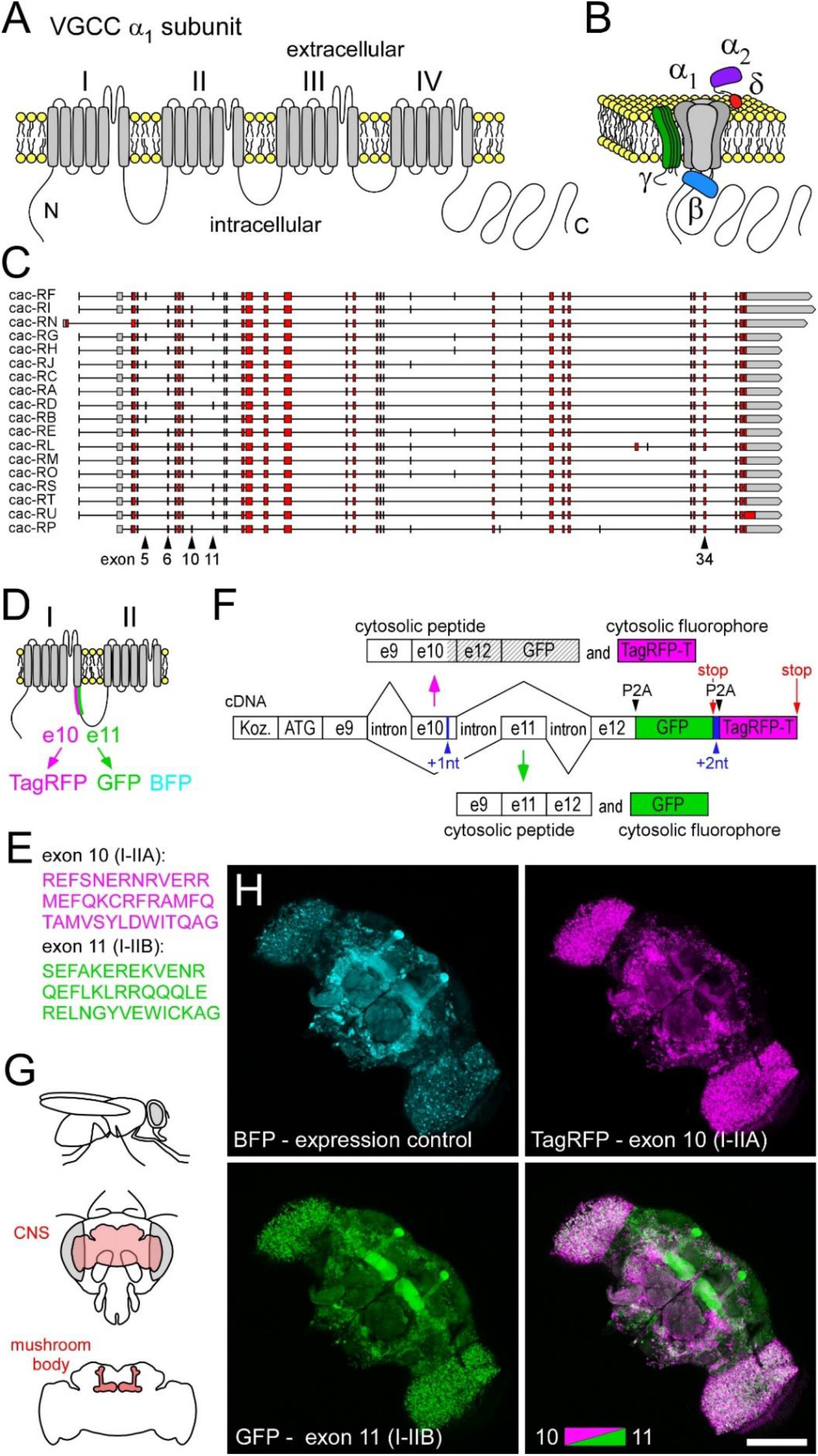
Bichromatic exon-reporters reveal biases in *cacophony* exon splicing between *Drosophila* neurons. A. A linearized representation of the voltage-gated Ca^2+^-channel (VGCC) α1-pore-forming subunit (polypeptide) Cacophony in the plasma membrane. Homologous domains I through IV are indicated, each containing 6 transmembrane segments. B. A 3D representation of the α1-subunit with accessory subunits β, α_2_δ and γ. C. Open reading frame representations of *cacophony* splice isoforms. Two pairs of mutually-exclusive exons (5 and 6, and 10 and 11) are indicated, along with the location of an exon (34) that is missing from only one of the 18 isoforms. D. Polypeptide context for location of mutually-exclusive exons 10 and 11 and their associated fluorophores, along with a third fluorophore (TagBFP) used as an independent expression control. E. AA sequences determined by exon 10 (I-IIA) and 11 (I-IIB). F. Schematic of DNA elements within a transgenic bichromatic exon-reporter, used to examine splicing bias of exons 10 and 11. Nucleotides are added (shown in blue) to move exons and fluorophores [GFP and TagRFP-T (TagRFP)] into frame, or out-of-frame, for subsequent translation, depending on pre-mRNA splicing. The vertical magenta and green arrows represent splicing (in the nucleus) followed by translation (in the cytosol) to yield cytosolic peptides. Hatching indicates out-of-frame peptides. G. Diagram of the orientation of the mushroom body within the brain, and thus the head capsule, of the adult fruit fly. H. A maximum projection of a series of confocal sections (7 images, each axially separated from the next by 2 um) through the brain of an adult. The exon 10 versus 11 reporter construct and TagBFP driven by the nSyb-GAL4 pan-neuronal driver. The brain was dissected from the head capsule prior to imaging.

Single-cell RNA sequencing (scRNA-seq) is routinely used to provide a transcriptional profile of individual neurons dissociated from vertebrate and invertebrate tissues and it shows a capacity to resolve many neuron types (Davie et al., 2018; Zeisel et al., 2015). Each neuron type might be further distinguished according to differences in their *spliceome* (Arendt-Tranholm et al., 2024; Joglekar et al., 2024), but such resolution is rarely achieved. While highly informative, scRNA-seq suffers from a number of limitations, such as transcriptional changes in response to the process of dissociating cells (Li, 2021), the loss of circuit/tissue context, and the absence of subcellular information. scRNA patch-seq, where the nuclear and cytosolic material is removed from the soma, allows for transcriptional profiling of neurons in their tissue context (Shao et al., 2023), and it has been used to identify *cacophony* isoforms in Drosophila *in situ* (Jetti et al., 2023). However, scRNA patch-seq is prohibitively difficult and a terminal assay. Furthermore, it is impractical to assay many neurons in the same preparation and cannot provide subcellular information.

Here, we sought to establish a technique that would allow us to simultaneously probe exon bias in *cacophony* in multiple neurons *in vivo*. Bichromatic fluorescent exon reporters have been used to monitor splice isoform bias *in vivo* in *C elegans* (Kuroyanagi et al., 2006), *Drosophila* (Li & Millard, 2019), mouse (Takeuchi et al., 2010) and rat (Oltean et al., 2008), but they have not been used to investigate VGCCs. The reporters previously used in *Drosophila* relied on modification of the gene’s endogenous locus (Gu et al., 2019; Li & Millard, 2019; Tadros et al., 2016) but as the *cacophony* locus barely tolerates modification (Gratz et al., 2019), we developed a transgenic approach. This approach allowed us to map the relative distribution of *cacophony* exons *in vivo* across groups of motor and sensory neurons in the larval periphery and across mushroom body structures in the adult brain. The ability to calibrate the reporter signal, and the consistency in stereotypical signal ratios across identified sets of neurons, demonstrates the utility of bichromatic fluorescent exon-reporters for tracking gene splicing in *Drosophila*, and for discovering and probing the neuron-specific function of previously undistinguished protein motifs in a VGCC α_1_-subunit.

## Results

### A bichromatic exon-reporter reveals splicing bias between *Drosophila* neurons

We initially probed splicing of exons 10 and 11 of *cacophony*, mutually-exclusive exons which code for alternate peptides that form the 5’ extent (40AAs) of the intracellular loop linking homologous domains I and II of the α_1_-subunit [I-IIA and I-IIB, respectively; http://flybase.org/ (Bell et al., 2024; Peixoto et al., 1997)] (Fig. 1D-E). Bichromatic exon-reporters are composed of DNA that mimic a limited stretch of a gene’s open reading frame (ORF) with the exons of interest and their bracketing introns, and DNA sequences encoding two distinct fluorophores (Fig. 1F). DNA of the reporter was integrated into the fruit fly’s second chromosome, and a GAL4 responsive Upstream Activation Sequence (UAS) allowed for its conditional expression. Spliceosomes will process pre-mRNA transcribed from both the gene’s endogenous locus and the reporter transgene. If the spliceosome retains exon 10, but excises exon 11, this construct will express TagRFP-T (TagRFP; Fig. 1F). Retaining exon 11 while excising exon 10, will result in EGFP (GFP) expression. An essential element of this design is to place GFP and TagRFP in different “frames”, and frame is then determined by pre-mRNA splicing. This design will also yield peptides corresponding to exons and fluorophores translated both in frame and out of frame. The potential for deleterious effects arising from competitive peptides is considered in a later section. TagBFP expression from a standard UAS construct was used as an expression control.

In an initial test of the utility of this construct we drove expression pan-neuronally and observed stark differences in fluorescence intensities of the two fluorophores in the brain of adult flies (Fig.1G-H). GFP fluorescence was particularly bright relative to TagRFP fluorescence in the mushroom body (lobes investigated in figure 3), while TagRFP was particularly bright in neurons within the optic lobes. Our interpretation was that this pattern indicated differences in the splicing bias between neurons in the MB and neurons in the optic lobes.

### Reporter expression levels vary across tissue types

To better interpret fluorophore expression patterns, we examined fluorescence intensities at the level of individual cells, which are readily identifiable at the larval neuromuscular junction (NMJ) (Fig. 2A). Established GAL4 drivers were used to drive the reporter construct for exon 10 versus 11 in motor neurons (MNs; nSyb), muscle (24B) and glia (Repo) (Fig. 2B-E). Each of three glutamatergic MNs form a terminal with big boutons [type-Ib (big)]; MN6/7-Ib forms a neuromuscular junction (NMJ) across muscle fibers #7 and #6, MN13-Ib forms a NMJ on fiber #13, and MN12-Ib forms a NMJ on fiber #12. A fourth MN (MNSNb/d-Is) forms a NMJ with small bouton terminals [type-Is, (small)] on each of muscle fibers #7, #6, #13 and #12 (in addition to others). *In vivo* examination revealed a higher level of GFP than TagRFP, relative to the expression control (TagBFP), in all MN terminals, indicating a splicing bias in favor of exon 11 (I-IIB) (Fig. 2F and G). Interestingly, there were significant differences in the ratio of GFP to TagRFP fluorescence between type-Ib axon terminals that belong to different MNs (Fig. 2H). However, the ratio of GFP to TagRFP fluorescence was no different between type-Is terminals that all issue from the axon of a single MN (MNSNb/d-Is). Construct expression in muscle fibers resulted in low-levels of GFP and TagRFP relative to the expression control (Fig. 2D, F-G), suggesting a low level of *cacophony* pre-mRNA splicing and functional VGCCs, as functional VGCCs require either exon 10 or exon 11. Expression in glia resulted in weak GFP expression but very strong TagRFP expression (Fig. 2E, G-I), suggesting that Repo positive perineural glia may use *cacophony* VGCC isoforms that rely on exon 10.

**Figure 2:**
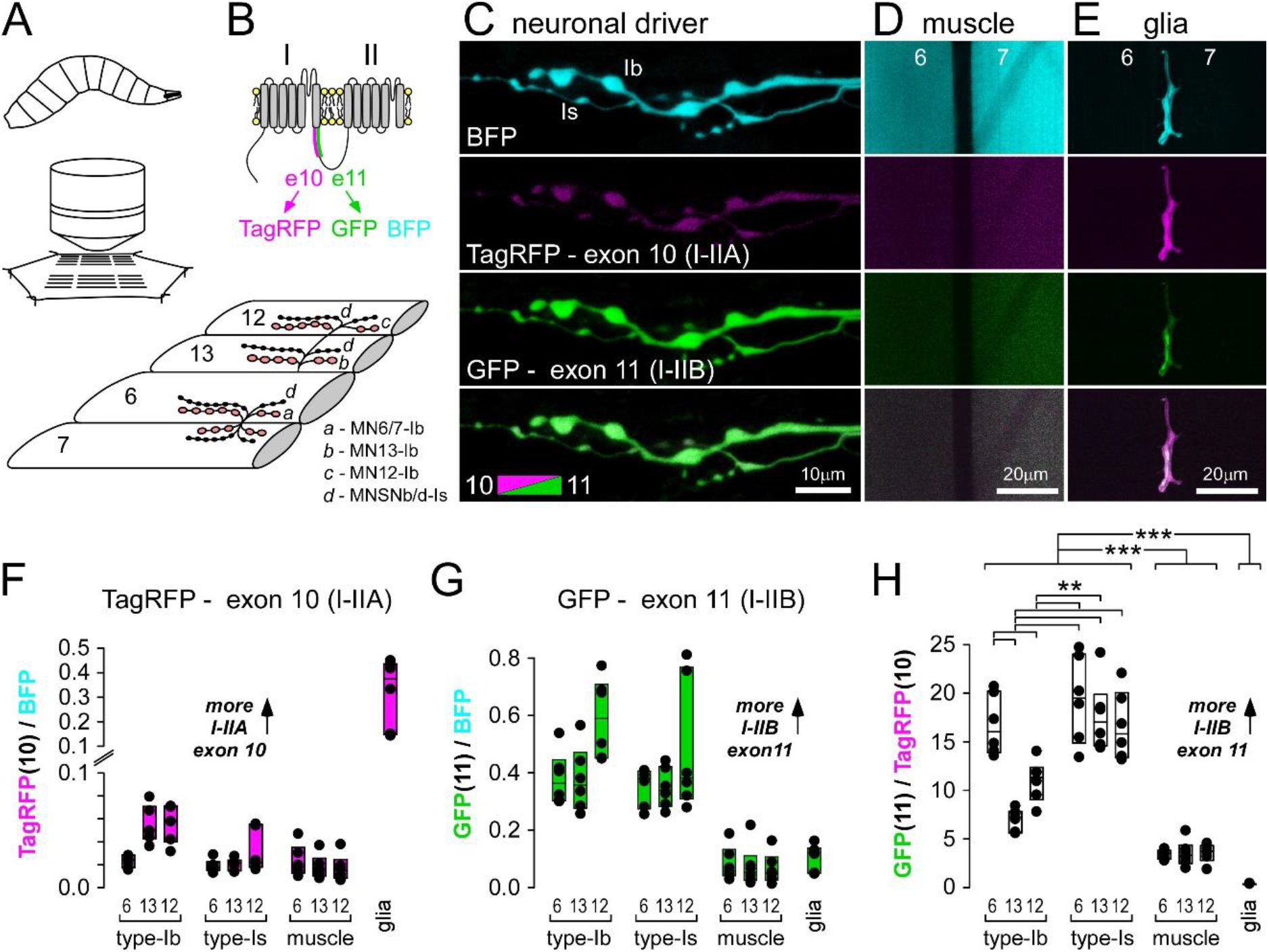
Differential splicing of *cacophony* exon 10 versus exon 11 across tissue types. A. Diagram of the general orientation of a larval filet dissection for live microscopic examination of neuromuscular junctions (NMJs) and perineural glia on identified body-wall muscle fibers #7, #6, #13 and #12. B. Polypeptide context for location of mutually exclusive exons 10 and 11, along with their associated fluorophores, and a TagBFP expression control. C. Presynaptic terminals of MN6/7-Ib and MNSNb/d-Is on muscle fiber #6, showing different levels of TagRFP and GFP expression relative to TagBFP, interpreted as different biases in the inclusion of exon 10 relative to exon 11 in mRNA. nSyb-GAL4 driver. D. Muscle fibers #6 and #7, showing TagRFP and GFP expression relative to TagBFP. 24B-GAL4 driver. E. Perineural glia showing TagRFP and GFP expression relative to TagBFP, between muscle fibers #6 and #7. Repo-GAL4 driver. F. Fluorescence intensity of TagRFP relative to the TagBFP expression control in different nerve terminals, muscle fibers and glia. Each point represents a different larval preparation. G. Fluorescence intensity of GFP relative to the TagBFP expression control. H. The ratio of GFP to TagRFP fluorescence intensity. A one-way ANOVA shows significant differences between axon terminals using Holm-Sidak post-hoc tests (**: P<0.005), while a second one-way ANOVA on ranks and post-hoc Dunn’s Method shows differences between tissue types (***; P<0.001). Boxes represent 25%-75%, while the line marks the median.

### The reporters reveal differences between mushroom body structures and peripheral sensory neurons

The reporter construct for exon 10 versus 11 also revealed stereotypical and highly consistent reporter ratios between lobes of the adult MB, and peripheral sensory neurons of larvae (Fig. 3). The MB structure has been annotated according to the regions where cholinergic Kenyon cells (KCs) of the calyx extend their axons and it is divided into a number of overlapping lobes (α, α’, β, β’ and γ) (Crittenden et al., 1998) (Fig. 3A). *In vivo* examination revealed α’ and β’ lobes to be characterized by TagRFP fluorescence, representing exon 10 (I-IIA), while α, β and γ lobes were characterized by GFP fluorescence, representing exon 11 (I-IIB) (Fig. 3B-D). This pattern was highly consistent across multiple flies (Fig. 3E), suggesting that KCs contributing axons to α’ and β’ lobes primarily rely on VGCC α1-subunit isoforms using exon 10, represented by cac-RA, -RB, - RE, -RH, -RI, -RL, -RM, -RN, -RO and -RP, while cells contributing axons to α, β and γ lobes use exon 11, represented by cac-RC, -RD, -RF, -RG, -RJ, -RS, -RT and -RU.

**Figure 3:**
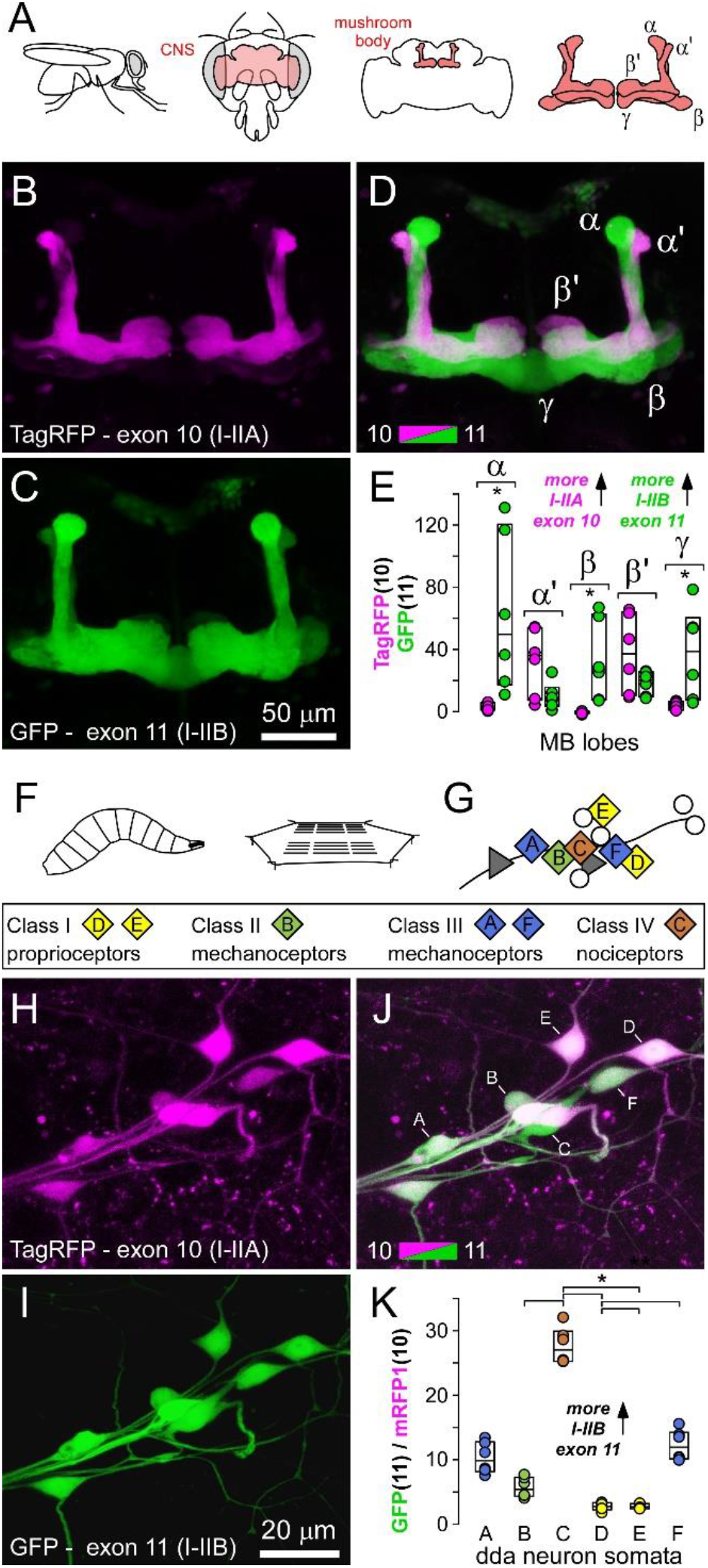
Stereotypical differences between mushroom body structures and sensory neurons. A. Diagram of the orientation of the mushroom body and its lobes, within the brain, and head capsule of the adult fruit fly. B-D. A mushroom body, with lobes identified, with the exon 11 (GFP) versus 10 (TagRFP) reporter construct driven by the nSyb-GAL4 pan-neuronal driver. E. The fluorescence intensity of GFP and TagRFP in the different lobes. Ratios were not plotted as faint lobular expression sometimes gave negative fluorescence values after background subtraction. Significant differences (P<0.01) in intensity are indicated with an asterisk (Mann Whitney Rank Sum tests; alpha adjusted to 0.01 after Bonferroni correction for 5 tests). F. Diagram of the general orientation of a larval filet dissection for live examination of dendritic arborization (da) neurons, sensory neurons that form multiple branched dendrites on the larval epidermis between the cuticle and body wall muscles. G. The somata of four different morphological classes of da neurons can be stereotypically identified in a dorsal cluster in each abdominal hemisegment [after Grueber and others (Grueber et al., 2003)]; ddaA through ddaF as shown. H-J. A dorsal cluster of sensory neuron somata with the exon 10 (TagRFP) versus exon 11 (GFP) reporter construct driven by the nSyb-GAL4 pan-neuronal driver. K. The ratio of GFP to TagRFP fluorescence intensity. A one-way ANOVA on ranks shows significant differences between axon terminals using Tukey pairwise post-hoc tests (asterisk: P<0.001).

The same reporter construct was examined in a dorsal cluster of dendritic arborization (da) neurons. These sensory neurons form multiple branched dendrites on the basal surface of the epidermis between the cuticle and body wall muscles (Fig, 3F-K). The somata of six da neurons, representing four different morphological classes can be stereotypically identified in a dorsal cluster in each abdominal hemisegment (ddaA-F; Fig. 3G) (Grueber et al., 2002). Here, we again observed strong stereotypical differences in fluorophore expression, and presumably exon bias (Fig. 3H-K) where exon 11 (I-IIB) is favored in all da neurons. Class category (I, II, III and IV) increases with increasing territory size and/or branching complexity (Grueber et al., 2002), and intriguingly, the ratio of exon 11 to exon 10 also increases with class category; Class I (D and E) < Class II (B) < Class III (A and F) < Class IV (C) (Fig. 3K).

### Reporting of splicing bias is independent of fluorophore choice

To determine whether differences in the ratio of GFP to TagRFP might be a function of different rates of protein maturation or degradation in different neurons, rather than differences in splicing *per se*, we reversed the order of fluorophores in the construct and quantified the ratios (Fig. 4). Unfortunately, TagRFP, which rarely forms aggregates, could not be used as it could not be placed 1 or 2 nucleotides out-of-frame without the creation of premature stop codons. Since mRFP1 does not generate stop codons when out of frame, it was paired with GFP to report the presence of exon 10 versus exon 11 (Fig. 4A-D), and vice versa (Fig. 4E-H). With mRFP1 in place of TagRFP the reporter construct duplicated the pattern of “*exon bias*” reported by the original TagRFP/GFP combination (compare Fig. 4D with Fig. 2H). Furthermore, the order of the fluorophores in the construct made little difference to the estimated ratio of exon 10 versus 11 when using the same microscopy settings for the different transgenes (Fig. 4I), i.e. the ratios in figure 4H show a reciprocal pattern to those in figure 4D.

**Figure 4:**
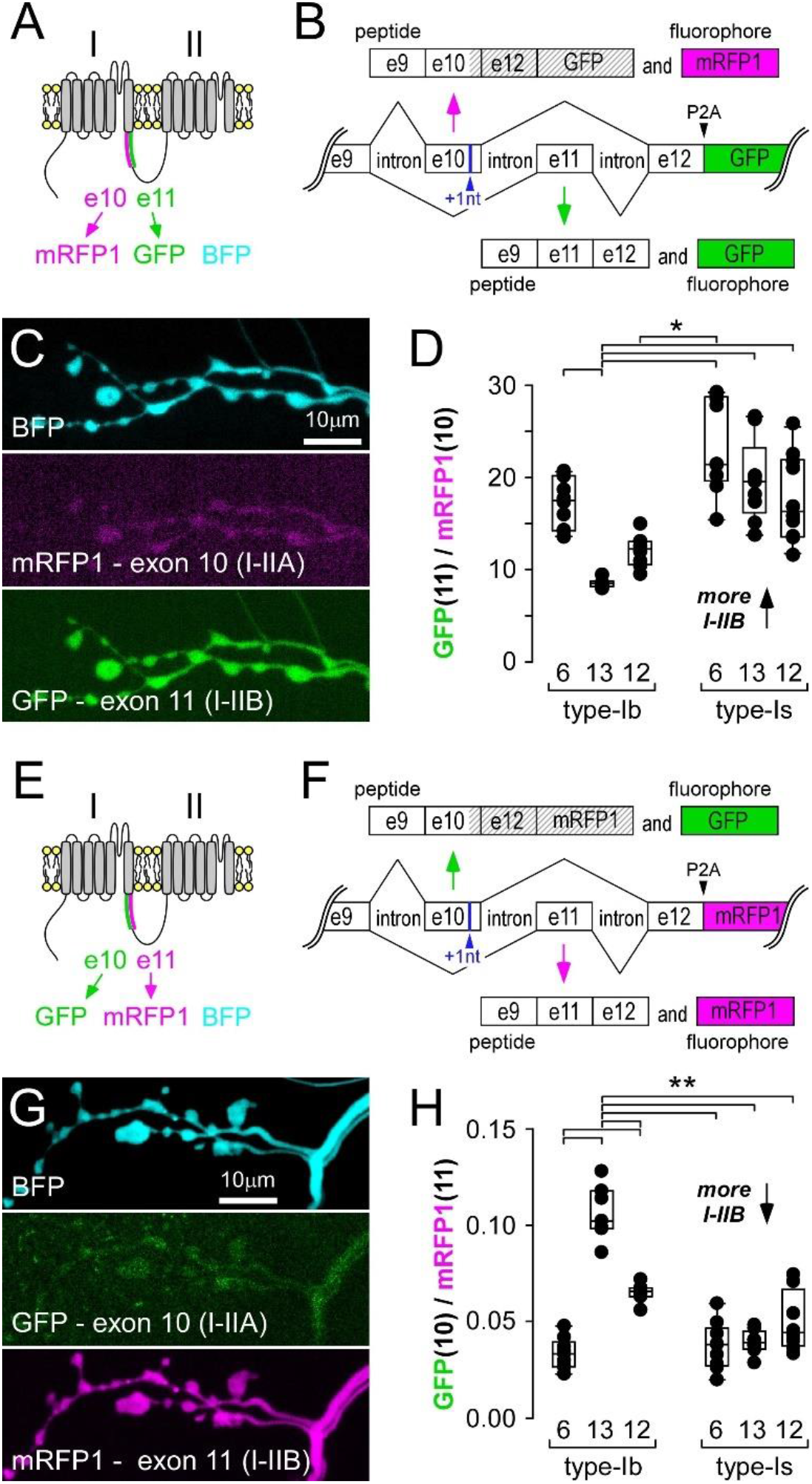
Measurements to calibrate bichromatic fluorescent exon-reporters. A. Protein context for location of mutually exclusive exons 10 and 11, their associated fluorophores, and a TagBFP expression control. B. Schematic of DNA elements within the transgenic bichromatic exon-reporter used to examine splicing bias of exon 10 versus 11. C. Presynaptic terminals of MN6/7-Ib and MNSNb/d-Is on muscle fiber #6 expressing the construct in B, and TagBFP, driven by nSyb-GAL4. D. The ratio of GFP to mRFP1 fluorescence intensity. A one-way ANOVA shows significant differences between axon terminals using Holm-Sidak post-hoc tests (asterisk: P<0.005). E and F. As in A and B, respectively, but with exon fluorophores switched. G. Presynaptic terminals of MN6/7-Ib and MNSNb/d-Is expressing the construct in F, driven by nSyb-GAL4. H. The ratio of GFP to RFP1 fluorescence intensity. A one-way ANOVA shows significant differences between axon terminals using Holm-Sidak post-hoc tests (**: P<gh240.001).

### Bichromatic exon-reporters can be calibrated

Reversal of fluorophore order also provides an opportunity to calibrate the ratio of fluorescence intensities in terms of a molar ratio, i.e. the relative number of fluorescent proteins. Reversal of the fluorophores allows us to nullify the effect of differences in microscope settings used for the different fluorophores. The fluorescence intensity *f* measured in each experiment can be expressed as:

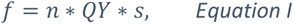

where n is the number of fluorophore proteins corresponding to the exon it represents, *QY* is the quantum yield of the fluorophore, and *s* represents the effect of microscope settings particular to that fluorophore, such as collection efficiency (depends on the numerical aperture of the objective lens and other optical elements), power of the excitation laser, gain of the photomultiplier, and extinction coefficient of GFP or mRFP1 at the excitation wavelength.

Using the Exon 10 GFP and Exon 11 mRFP1 construct, we can write:

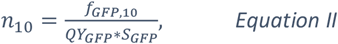

If we reverse the fluorophores such that mRFP1 reports exon 10, and GFP reports Exon 11 we can similarly write:

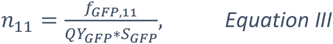

Using equations II and III, we can write:

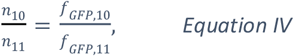

Similarly, we can obtain:

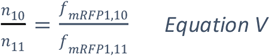

Alternatively, the ratio of proteins can be calculated using equation VI:

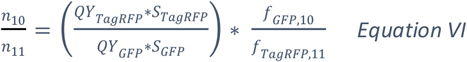

From equation VI, we can calculate the ratio of protein numbers, 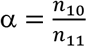, and the value of 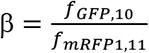 can be also determined experimentally. The unknown factor 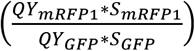 appearing in equation VI can be calculated using equation VII:

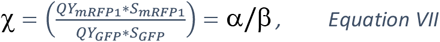

For the fluorophores we selected and microscope settings used, χ = 1.05. Knowing this value, the molar ratio of fluorophores representing the splice bias can be calculated by multiplying the ratio of their fluorescence intensities by χ, or

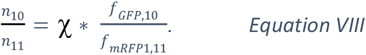

Our conclusion is that, through serendipity alone, the relative values of fluorescence intensities closely represent the relative numbers of fluorophores.

### The reporters reveal consistent differences in splicing bias of other exons

Bichromatic exon reporters can be used to interrogate splicing of other mutually exclusive exons (5 versus 6), as well as a seemingly non-essential exon (34) (Lembke et al., 2019) (Fig. 5). Exons 5 and 6, each determine the identity of two different peptides of 35AAs which form the S4 voltage sensor in the first homologous domain [IS4A and IS4B, respectively; http://flybase.org/ (Bell et al., 2024; Peixoto et al., 1997)] (Fig. 5A-B). The ratio of fluorophores corresponding to exon 5 (mRFP1) versus exon 6 (GFP) (Fig. 5C) was consistent between MNs from one larva to the next (Fig. 5D-E), just as it was with mutually exclusive exon 10 versus 11 (Figs. 2 and 4). The ratios suggest that motor neurons rely most heavily on VGCC α1-subunits using exon 6 (IS4B), an exon that carries less positive charge in the S4 voltage sensor compared to the alternative exon (5; IS4A).

**Figure 5:**
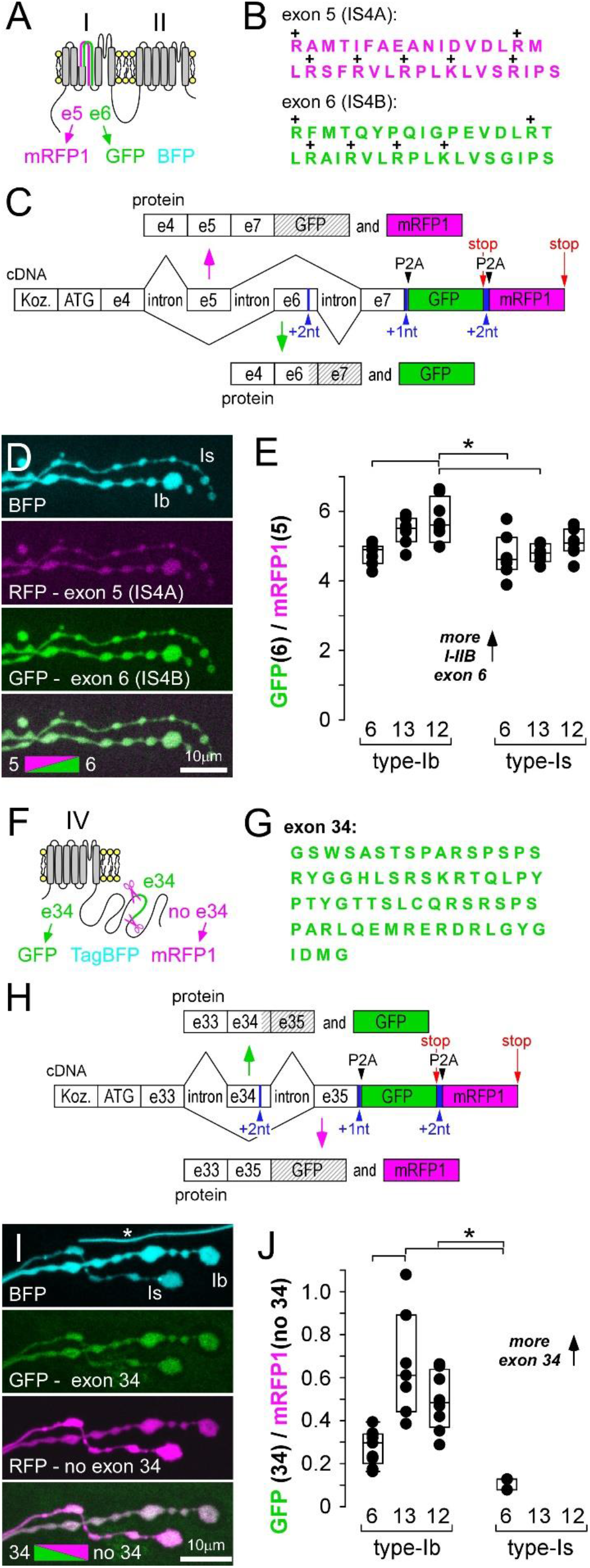
Splice reporters for exons 5 versus 6, and 34, also show stereotypical patterns of bias. A. Protein context for location of mutually exclusive exons 5 and 6, their associated fluorophores, and a TagBFP expression control. B. AA sequences determined by exons 5 and 6. C. Schematic of DNA elements within a transgenic bichromatic exon-reporter, used to examine splicing bias of exons 5 and 6. D. Presynaptic terminals of MN6/7-Ib and MNSNb/d-Is on muscle fiber #6, showing mRFP1 and GFP expression relative to TagBFP. nSyb-GAL4 driver. E. The ratio of GFP to mRFP1 fluorescence intensity. A one-way ANOVA shows significant differences between axon terminals using Holm-Sidak post-hoc tests (*: P<g245.0.005). F. Protein context for location of exon 34, the associated fluorophores, and a TagBFP expression control. G. AA sequences determined by exon 34. H. Schematic of DNA elements within a transgenic bichromatic exon-reporter, used to examine splicing bias of exon 34. I. Presynaptic terminals of MN6/7-Ib and MNSNb/d-Is on muscle fiber #6, showing GFP and mRFP1 expression relative to TagBFP. Asterisk denotes autofluorescence from a fine tracheole. nSyb-GAL4 driver. J. The ratio of GFP (includes exon 34) to mRFP1 (exon 34 excised) fluorescence intensity. A one-way ANOVA shows significant differences between axon terminals using Holm-Sidak post-hoc tests (*: P<0.005). MNSNb/d-Is terminals were missing from muscles fibers #13 and #12.

Exon 34 determines the identity of 68AAs in the middle of the carboxy terminus in all but one Cacophony isoform (cac-RM) [http://flybase.org/ (Chang et al., 2014; Lembke et al., 2019; Lembke et al., 2017)] (Fig. 1C, 5F-G). Surprisingly, we found that the fluorophore corresponding to splicing exon 34 from the pre-mRNA (mRFP1), was present in all neurons, and present at higher levels than the fluorophore corresponding to the presence of exon 34 (GFP), indicating high levels of the cac-RM isoform in all cells. The ratio reported in MNSNb/d-Is terminals indicated that these terminals had the lowest proportion of Cacophony isoforms with exon 34, but the same terminals were often physically missing.

### The reporters reveal differences between neurons in the larval ventral ganglion

To determine the extent to which larval central neurons might manifest differences in exon bias, we drove expression with a pan-neuronal driver (nSyb) and examined patterns within the ventral ganglion (Fig 6). *In vivo* examination revealed diverse but highly consistent patterns from one larva to the next. The consistency is demonstrated by the common patterns observed across different larvae for each reporter construct (Fig. 6B, D, G, J, M). The patterns were consistent when different fluorophores were used, i.e. mRFP1 used in place of TagRFP for mutually exclusive exons 10 and 11 (compare solid line inset of Fig. 6C with 6E), and the pattern inverted when the fluorophores were reversed (compare Fig. 6E with 6H).

**Figure 6:**
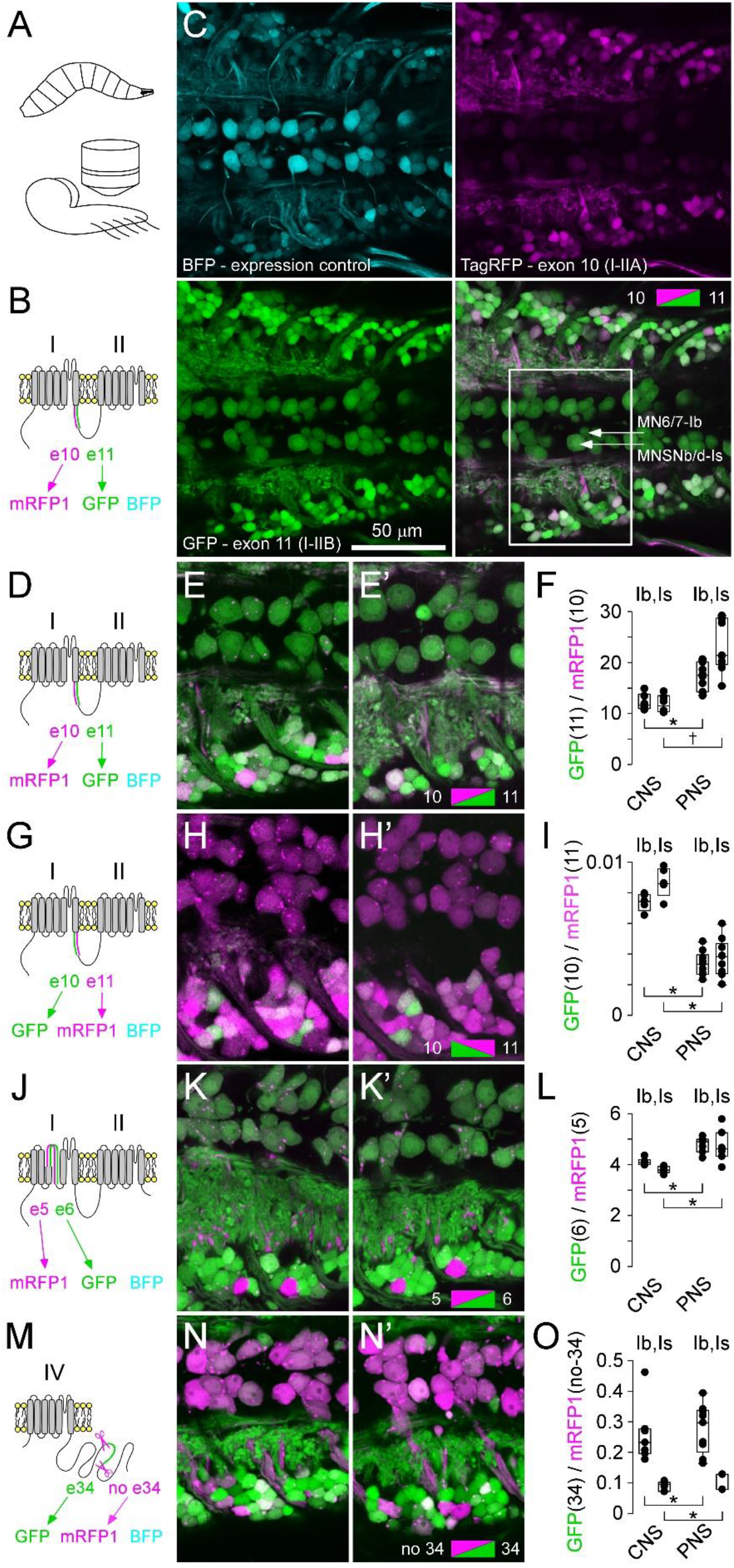
Stereotypical patterns of exon bias in the larval ventral ganglion. A. Diagram of the general orientation of a larval ventral ganglion preparation for live microscopic examination. B. Polypeptide context for location of mutually exclusive exons 10 and 11, and a TagBFP expression control. C. Neurons in the larval ventral ganglion (VG) expressing the exon 10 (TagRFP) versus 11 (GFP) reporter construct (Figs. 1-3) and a TagBFP expression control. D, G, J, M. Polypeptide context for location of probed exons – data grouped in rows. E-E’, H-H’, K-K’, N-N’. Fields-of-view, similar in extent to the solid line inset in C, spanning hemisegments 3 and 4. Two separate larval VGs are shown in each of E-E’, H-H’, K-K’, N-N’. In each case, the nSyb-GAL4 driver was used and a single confocal section captured most of the dorsal midline MN somata. F, I, L, O. The fluorophore ratio intensity measurements for each reporter shown immediately to the left, for somata of MN6/7-Ib and MNSNb/d-Is in the central nervous system (CNS), and their terminals in the periphery (PNS). CNS location of the somata shown in inset in C. Asterisks indicate significant differences (P<0.005) in Student’s t-tests. Alpha adjusted to 0.025 after Bonferroni correction for 2 tests. Dagger indicates a significant difference (P=0.002) in a Mann Whitney Rank Sum test.

### The reporters suggest subcellular differences in exon bias

To determine whether the pattern observed in the somata represents the same pattern observed in the axon terminals of the same neurons, we compared the peripheral values with the central values (Fig. 6F, I, L, O). Reporter ratios indicate a bias towards exon 11, relative to exon 10, in the terminals compared to somata in both terminal types (Fig. 6F). This was unexpected, as pre-mRNA splicing occurs in the nucleus (but see Discussion), and we anticipated a constant ratio across all parts of the neuron if GFP diffuses as readily as mRFP1. A P2A peptide between exon-associated peptides and the fluorophore ensures that the fluorophore is free to diffuse unencumbered by other peptides. When fluorophores were reversed, relative to the exons (Fig. 4F vs 4B), the significant bias towards exon 11 in the terminals was preserved (compare Fig. 6I with 6F). A small but significant bias towards exon 6, relative to exon 5, was also observed in terminals compared to their somata (Fig. 6L). No bias was observed towards exons containing exon 34 in terminals (Fig. 6O).

### *In vivo* presynaptic properties are consistent with exon reporter ratios

To determine whether the fluorophore ratios established here do indeed represent a splicing bias, and presumably the mRNA ratio of *cacophony* splice isoforms, we tested physiological predictions based on exon reporter ratios. We proposed that exon reporter ratios reflect VGCC peptide ratios and made predictions regarding presynaptic physiology based on biophysical and physiological data associated with the inclusion/exclusion of either exon 10 or 11 and exon 5 or 6 (Bell et al., 2024) and exon 34 (Chang et al., 2014; Lembke et al., 2019; Lembke et al., 2017). We then tested these predictions against our own terminal specific physiological data, both published (Chouhan et al., 2012; Chouhan et al., 2010; Justs et al., 2022; Lu et al., 2016) and unpublished.

As previously described (Lu et al., 2016), we have used electron micrographs from multiple sets of serial sections through each terminal type to make estimates of the average number of AZs at each MN terminal (Table 1). These data were coupled with our estimates of quantal content and previous Ca^2+^ entry data (Justs et al., 2022), to make estimates of the *average* probability of release at each AZ (P_AZ_; Fig. 7E, I, M) and *average* Ca^2+^ entry at each AZ (Fig. 7F, J, N) across all terminal types (Table 1). No suggestion is made that release probability is uniform across AZs in the same presynaptic terminal as non-uniformity is well documented at the *Drosophila* NMJ (Newman et al., 2022), but rather, P_AZ_ is terminal specific and a useful metric to compare with the proposed terminal-specific peptide ratios. Estimates of endogenous firing frequency particular to each terminal were taken from Chouhan and others (Chouhan et al., 2012; Chouhan et al., 2010) (Fig. 7G, K, O) (Table 1).

**Table 1:** Physiological data specific to each of the 6 different motor nerve terminals. These data represent new data or analyses, annotated as “present”, or a compilation from previous studies published by our laboratory annotated as “Justs” (Justs et al., 2022) or “Chouhan” (Chouhan et al., 2012; Chouhan et al., 2010). Whole terminal volume data (column 2) (μm^3^) were generated through immunohistochemistry and confocal microscopy, as described previously (Justs et al., 2022; Lu et al., 2016) and published in (Justs et al., 2022). AZ number per unit volume (μm^-3^) data (column 3) were generated through reference to transmission electron microscopy data described previously (Justs et al., 2022; Lu et al., 2016), but mostly unpublished. Total AZ number for specific terminals (column 4) is the mathematical product of columns 2 and 3. Quantal Content (QC) data were generated from terminal-specific TEVC recordings of EJCs, and muscle-specific TEVC recordings of mEJCs, all in the presence of 2mM CaCl2 added to HL6 (see Methods). The average probability of release from an individual AZ (P_AZ_; column 6) was calculated by dividing QC by the total number of AZs. The terminal-specific change in [Ca^2+^]_i_ in response to a single isolated AP (column 7) was reported previously (Justs et al., 2022), and was measured using forward-filled rhod dextran mixed with AF647. The number of Ca^2+^ ions entering each terminal (column 8), and the method of calculation was reported previously (Justs et al., 2022). The average number of Ca^2+^ ions entering through each AZ in response to an AP (column 9) is calculated by dividing total Ca^2+^ entry by the number of AZs. The endogenous firing frequency of each MN terminal (column 10) was reported previously (Chouhan et al., 2012); calculated through the use of current clamp electrodes recording simultaneously in adjacent muscles fibers in the presence of 2mM CaCl added to HL6. Standard error of the mean (SEM) calculated according to propagation of uncertainty theory (Farrance & Frenkel, 2012) in columns 4, 6 and 9.

| terminal identity | volume<br>($\mu\text{m}^3$)<br>*Justs* | AZs/vol.<br>($\mu\text{m}^{-3}$)<br>*present* | total AZs<br>($N_{\text{AZ}}$)<br>*present* | Q.C.<br>(quanta)<br>*present* | AZ prob.<br>($P_{\text{AZ}}$)<br>*present* | $\Delta[\text{Ca}^{2+}]_i/\text{AP}$<br>(nM)<br>*Justs* | $\text{Ca}^{2+}/\text{AP}$<br>($10^6$ ions)<br>*present* | $\text{Ca}^{2+}/\text{AP}/\text{AZ}$<br>($10^3$ ions)<br>*present* | firing<br>(Hz)<br>*Chouhan* |
| --- | --- | --- | --- | --- | --- | --- | --- | --- | --- |
| MN6/7-Ib | 311±20 | 2.62±0.22 | 813±86 | 81.9±4.8 | **0.101±0.012** | 305±39 | 3.79±0.30 | **4.66±0.62** | **21.3±0.5** |
| MN13-Ib | 392±49 | 2.21±0.31 | 864±163 | 38.7±9.5 | **0.045±0.014** | 160±27 | 2.85±0.33 | **3.30±0.73** | **42.0±1.1** |
| MN12-Ib | 366±38 | 2.21±0.56 | 808±222 | 25.5±2.5 | **0.032±0.009** | 253±36 | 4.03±0.33 | **4.99±1.43** | **32.4±1.7** |
| MNSNb/d-Is, M6 | 90±20 | 2.69±0.36 | 242±63 | 74.2±3.8 | 0.306±0.081 | 512±109 | 1.67±0.29 | 6.89±2.15 | 7.8±0.6 |
| MNSNb/d-Is, M13 | 97±16 | 6.03±1.78 | 584±197 | 43.2±5.6 | 0.074±0.027 | 420±88 | 1.54±0.23 | 2.64±0.98 | — |
| MNSNb/d-Is, M12 | 128±29 | 4.13±0.71 | 530±151 | 62.3±5.6 | 0.117±0.035 | 522±86 | 2.43±0.36 | 4.59±1.47 | 7.8±0.1 |
| Is average |  |  |  |  | **0.166±0.071** |  |  | **4.71±1.23** | **7.8±0.3** |

**Figure 7:**
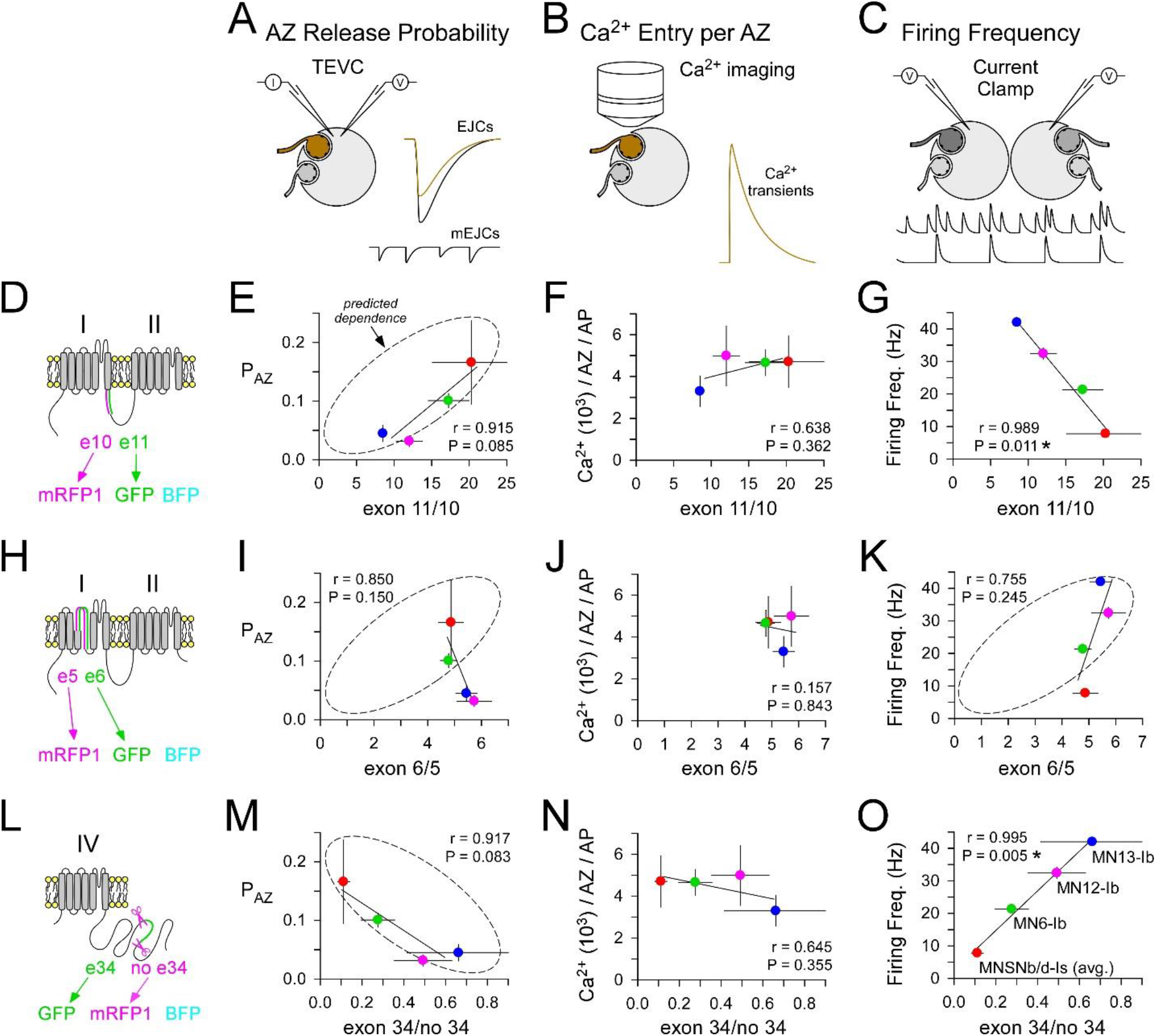
Physiological data validate several predictions from the reported exon biases. A-C. Stylized diagrams representing the techniques used to collect physiological data plotted in the corresponding columns below each technique. Terminal specific data in Table 1. A. Two-electrode voltage clamp (TEVC) used to estimate a terminal’s quantal content (QC). The average probability of release at individual active zones (P_AZ_) is calculated as the QC for a single isolated AP divided by the average number of AZs (Table 1, column 6). B. Ca^2+^ imaging with chemical Ca^2+^ indicators used to estimate the total number of calcium ions (Ca^2+^) that enter a terminal. The average number of Ca^2+^ ions (10^3^) entering each AZ during an AP (Ca^2+^_AZ_) is calculated as the total Ca^2+^ entry divided by the average number of AZs (Table 1, column 9). C. Current clamp recordings from adjacent muscle fibers used to estimate terminal specific endogenous firing frequencies (Table 1, column 10). D,H,L. Polypeptide context for the data in each adjacent row. E,F,G. Physiological data (legend in O) plotted against the exon 11 vs 10 ratio. Where a prediction has been made regarding the influence of the exon ratio an ellipse has been drawn with the long axis denoting whether a negative or positive dependence is predicted. I,J,K. Physiological data plotted against the exon 6 vs 5 ratio. M,N,O. Physiological data plotted against the exon 34 present to 34 absent ratio. Pearson product-moment correlation coefficient was calculated to test the strength and direction of associations. Least-squares linear fits. Legend for terminal identity is shown in O. Red: MNSNb/d-Is (average from three muscle fibers); Green: MN6-Ib; Pink: MN12-Ib; Blue: MN13-Ib.

Bell and others (Bell et al., 2024) established that alternate splicing of exon 10 versus 11 affects Cacophony number and release probability. Specifically, excision of exon 11 (I-IIB), and not exon 10 (I-IIA), results in a reduction in neurotransmitter release and less Cacophony at each AZ. We would therefore expect a positive correlation between the ratio of exon 11 relative to exon 10 and P_AZ_, and this is what we observed (r=0.915; P=0.085) (Fig. 7E), and is generally consistent with our estimates of Ca^2+^ entry at each AZ across terminals (r=0.638; P=0.362) (Fig. 7F). While no predictions are made regarding firing frequency (but a prediction is made with regard to exon 5/6) we do see a significant negative association between firing frequency and the ratio of exon 11 to 10 (r=0.989; P=0.011) (Fig. 7G).

Bell and others (Bell et al., 2024) established that exon 6 (IS4B) is required for Cacophony localization to the active zone and tuning Cacophony’s biophysical properties, and that exclusion of exon 6 from the *cacophony* locus (but not exon 5; IS4A) is embryonic lethal. Consistently, exon reporters show exon 6 to be present at high levels in all terminals. Of all the exons examined, the ratio of exons 6 to exon 5 showed the least variation, thus providing little leverage for testing predictions. Based on the data of Bell and others, we might expect to see higher P_AZ_ with a higher level of exon 6, but this is not seen (Fig. 7I). Bell and others found that the presence of exon 6 is required to prevent inactivation during sustained depolarization. Naranjo and others (Naranjo et al., 2015) found that zebrafish Ca_v_2.1 do not inactivate completely during sustained depolarization, a property compatible with carrying action potentials at higher rates. We might therefore expect that higher levels of exon 6 would be required to allow motor neurons to fire at high rates. Consistently, we found an increase in firing frequency with an increase in the ratio of exon 6 to exon 5 (r=0.755; P=0.245) (Fig. 7K).

Lembke and others (Lembke et al., 2019; Lembke et al., 2017) found no deficit in evoked release at the NMJ when exon 34 was excised from the endogenous locus, indicating that the sole cac-RM isoform is competent to traffic to the plasma-membrane and mediate AP-triggered neurotransmitter release. Furthermore, under these conditions they reported Cacophony levels (the protein) to be diminished to 20% of wildtype levels, despite mRNA levels being undiminished. The implication is that, Cacophony, in the *absence* of exon 34, is a potent facilitator of neurotransmitter release. Therefore, we predict that higher levels of exon 34 would result in *less* neurotransmitter release, and this is what we observe with a negative correlation between P_AZ_ and the reported levels of exon 34 (r=0.917; P=0.083) (Fig. 7M). The published data cannot be easily be extrapolated to make predictions regarding either Ca^2+^ entry or endogenous firing frequencies (Fig. 7N and O; see Discussion). In conclusion, where we find a substantial spread in exon reporter ratios between neurons (11 vs 10, and 34 vs no-34), we also find substantial spread in terminal specific physiological properties that might be predicted if exon reporter ratios reflect peptide ratios.

## Discussion

Here, we investigated alternative splicing in *cacophony* across multiple neurons *in vivo* using fluorescent bichromatic exon reporters. These reporters indicated a wide range of exon bias between neurons, providing insight into the role of alternate peptides in polypeptide function and circuit context. Furthermore, we revealed subcellular differences, information that might only otherwise be gleaned using isoform-specific antibodies or genetic tags (should they be available). As the pattern of exon bias is highly consistent from one animal to the next it suggests that each neuron splices a consistent and unique portfolio of VGCC isoforms. However, these reporters are only valuable if it can be established that they do indeed report splicing biases. Serendipitously, each exon has been investigated for its physiological relevance in *Drosophila*, allowing us to make testable predictions for a set of motor neurons innervating body wall muscles. These predictions were generally consistent with our Ca^2+^ entry data, release site probability data and endogenous firing frequency data, indicating that our exon reporters do reveal *cacophony* exon splicing biases, if not peptide biases.

While *cacophony* is homologous to α_1_-subunit genes within the vertebrate Ca_v_2 VGCC family, the exons studied here appear to be without homology in vertebrate Ca_v_2 α_1_-subunits, yet they appear to serve a crucial role in providing functional diversity in *Drosophila*. In vertebrates, different Ca_v_2 α_1_-subunits contribute differently to the strength and plasticity of neurotransmitter release (Catterall & Few, 2008; Regehr & Mintz, 1994) as do the different splice isoforms of these Ca_v_2 α_1_-subunits (Bourinet et al., 1999; Cingolani et al., 2023). In *Drosophila*, alternative splicing of *cacophony*’s many exons allows for functional specialization across different neuronal circuits, compensating for the limited number of Ca_v_2 α_1_-subunit genes. Thus, despite the lack of direct homologues, the study of these exons in *Drosophila* can offer valuable insights into how alternative splicing can generate a diverse range of VGCC isoforms and functions, with potential implications for understanding similar processes in more complex vertebrate systems.

A simplistic model of the mode of action of these reporter constructs is that cells fill with a mixture of fluorophores reflecting the composition of the pool of mRNA immediately outside the nucleus, and that this reflects spliceosome bias. However, this is an unrealistic model as transcription rates change along with spliceosome bias throughout the lifetime of a neuron (Furlanis & Scheiffele, 2018). Such changes will evade detection by these reporter constructs as their signal is “integrative” in nature, providing a long-lasting trace of spliceosome activity. The constructs might be engineered to show greater temporal resolution through the inclusion of degradation signals, a strategy that would trade signal strength for temporal responsiveness. Alternatively, construct transcription might be conditionally activated using techniques such as GeneSwitch or Gal80^ts^ (McGuire et al., 2003; Osterwalder et al., 2001). A more acute approach might be to photobleach or photoconvert the fluorophores, allowing for quantification of newly translated fluorophore.

Our observation of differences in the fluorophore ratio between soma and axon terminals in the very same neurons (Fig. 6F,I,L) highlights the limitations of a reporter design that relies on translation of mRNA to signal spliceosome bias, but potentially reveals an interesting phenomenon. If fluorophores simply fill the extremities through diffusion or transport of protein from ribosomes in the soma, we would have expected a similar ratio in the terminals as in the somata. The P2A peptide between exon-derived peptides and the fluorophore ensures that the fluorophores travel alone. Axon terminals are translationally competent in *Drosophila* as in other organisms (Holt et al., 2019; Nesler et al., 2016), but if transcripts are trafficked to neuronal extremities without isoform specificity, translation alone seems unlikely to give rise to fluorophore ratio differences between terminals and somata. However, if cytoplasmic splicing occurs within these axon terminals, as shown in mammals (Nikolaou et al., 2022; Salehi et al., 2023), then such a mechanism would allow for a change in the fluorophore ratio at peripheral ribosomes. As transcripts are actively transported (Das et al., 2021), an alternative explanation is that mRNA isoforms are being differentially transported into the terminals; the outcome of multiple overlapping mechanisms (Gao et al., 2008). If this is what our data have revealed in *Drosophila* larval MNs, transcripts containing exon 11 (I-IIB) appear to be twice as likely as those with exon 10 (I-IIA) to be transported into axon terminals, and transcripts containing exon 6 (IS4B) are transported with a marginally higher probability than those with exon 5 (IS4A). Our data are mute on the possible differential transport of VGCC α_1_-subunit splice isoforms (the endogenous protein) to different destinations within the neuron, but the data here suggest that a similar result might be achieved by differential transport of mRNAs. Spatial resolution of these reporters might be improved using the strategies described above; the same strategies that improve the temporal resolution of the reporter signal.

While we made predictions regarding average Ca^2+^ entry at individual AZs based on published neurotransmission data (Bell et al., 2024; Lembke et al., 2019), we could not make predictions regarding average VGCC number at individual AZs. Such an extrapolation is not sustainable as neither average Ca^2+^ entry nor the average probability of release from an AZ (P_AZ_) is positively correlated with average VGCC number between AZs at different *Drosophila* nerve terminals (Medeiros et al., 2024). For example, while Ca^2+^ entry at the AZs of MNSNb/d-Is terminals is greater than that at the AZs of MN6-Ib (He et al., 2023; Lu et al., 2016) and the probability of release is between 2- and 3-fold higher at the AZs of MNSNb/d-Is (Lu et al., 2016; Newman et al., 2022), VGCC number is no different (He et al., 2023; Medeiros et al., 2024). In other words, factors apart from VGCC number play a major role in determining Ca^2+^ entry at different terminals, e.g. differences in VGCC splice isoforms, accessory subunits, post-translational modifications or even differences in AP width. Without identifying terminals with different numbers of Cacophony VGCCs in their AZs, our information on exon bias between terminals cannot be leveraged to determine which exons might influence Cacophony VGCC number. However, we can still identify our best exon candidates by identifying those exons which have the same ratio in terminals where AZs have the same number of VGCCs. In this case, exon 6 versus 5, or exon 11 versus 10, would be our best candidates as they show no difference in ratio between MNSNb/d-Is and MN6-Ib terminals (Fig. 7E and 7I); terminals with no difference in VGCC number at their AZs (He et al., 2023; Medeiros et al., 2024).

Exon 34 is missing from only 1 of the 18 annotated cacophony isoforms, and yet this isoform (Cac-RM) is present at only low levels in RNA-seq data quantification measures performed by Jetti and others (Jetti et al., 2023). Jetti and others performed patch-seq RNA profiling on the material extracted from the somata of identified motor neurons in the 3^rd^ instar larva, and reported the gene expression measure of total transcripts per million (TPMs). They reported that mRNA for the Cac-RM isoform was present at only 12.9% of total TPMs in a type-Ib representative (MN1-Ib) and only 11.4% in a type-Is representative (MNISN-Is). In contrast, our data, also from the 3^rd^ instar larva, indicate that there is a bias towards the exclusion of exon 34, i.e. only 30-40% of isoforms include exon 34. Surprisingly, Lembke and others (Lembke et al., 2019) deleted exon 34 in the genome and found that it (and the 17 isoforms that it appears in) is not required for *Drosophila* viability. However, these 17 isoforms are by no means redundant, as exon 34 deletion results in a disrupted motor pattern output and slower locomotion in larvae. Yet, despite an 80% reduction in Cacophony protein measured in the adult nervous system, they did not find a deficit in evoked neurotransmitter release at the larval NMJ.

Our reporters indicated that MNSNb/d-Is terminals, which have the highest level of Ca^2+^ entry per AZ (He et al., 2023; Lu et al., 2016), had the lowest proportion of Cacophony isoforms with exon 34. These terminals were often missing from muscle fibers #6 and #7, and always missing from #13 and #12. Missing terminals are a rare phenomenon and may represent a developmental phenotype arising from the presence of competitive peptides; either in-frame or out-of frame. Out-of-frame fluorophore peptides are common to all of these reporters and so it is unlikely that they are responsible for the phenotype. Along with high levels of mRFP1, MNSNb/d-Is will contain high levels of in-frame exons 33 and 35 peptides (Fig. 5H) implicating roles for either peptide, or both, in the “missing terminals” phenotype. Adaptor protein complex-1 (AP-1) binds a site in the carboxy terminus of mouse Ca_v_2.2 and assists its trafficking to the plasma membrane (Macabuag & Dolphin, 2015). Lembke and others (Lembke et al., 2019) identified an AP-1 binding domain in the exon 35 peptide of Cacophony, and this raises the possibility that competitive peptide binding disrupts Cacophony trafficking to the plasma-membrane. The identity of exon 34 as a binding site for *tdph*, the *Drosophila* ortholog of transcription factor TAR RNA-binding protein (TDP-43), is intriguing, as are the phenotypic similarities between *tdph* mutants and exon 7 excision mutants (Chang et al., 2014; Hazelett et al., 2012; Lembke et al., 2019; Lembke et al., 2017), but the role of exon 34 in transcriptional regulation is far from clear. The 3’ end of exogenous exon 34 is translated out of frame (Fig. 5H) in type-Ib MNs, producing a peptide with further potential for a dominant negative role, but no developmental phenotypes were observed in type-Ib terminals.

Our observation of differential exon biases between MB lobes (Fig. 3B-E) suggests differential involvement of VGCC isoforms in olfactory information processing. Kenyon Cells (KCs) are intrinsic to the MB where they fasciculate as they course through the peduncles and terminate in separate branches of the dorsal and medial lobes. KCs that terminate in the γ lobe are associated with short term memory, while KCs of the α′ and β′ lobes play a role in memory consolidation and KCs of the α and β lobes are crucial for long-term memory (Davis, 2023). The preferential expression of exon 10 in the γ lobe and α′ and β′ lobes suggests that exon 10 (I-IIA) may play specific roles in both short-term memory and memory consolidation. Conversely, the preferential expression of exon 11 in α and β lobes suggests that exon 11 (I-IIB) may be necessary for the long-term synaptic changes required for memory storage. Therefore, a fine-grained map of exon bias across lobes of the MB might offer substantial insight into the contribution of VGCC α_1_-subunit splice isoform diversity to olfactory information processing. In conclusion, this study has established the utility of transgenic bichromatic exon reporters for investigating the role of alternative splicing in neuronal function in *Drosophila*. By mapping exon usage *in vivo* across different neuronal types, this highly adaptable and visually intuitive tool will provide another avenue for elucidating genetic and molecular mechanisms underlying neuronal diversity and circuit function.

## Methods

### Fly stocks

*Drosophila* stocks were raised at 24°C on standard medium [Bloomington Drosophila Stock Center (BDSC) recipe]. Measurements were performed on female 3^rd^ instar larvae of a w^1118^ isogenized strain. BDSC (Bloomington, IN) provided the following fly lines: 24B-GAL4 (stock #1767); Repo-GAL4 and nSyb-GAL4 (stock #51635). UAS-TagBFP was a gift from Dr Kenneth Irvine.

### Fluorescent Bichromatic Exon-Reporter Design

Each construct started with a TATA box sequence followed by the standard start codon (ATG). The full DNA sequence of the *Drosophila cac* gene was obtained via FAST sequence from NCBI which was then aligned with the cDNA sequences including or lacking exons of interest using MUltiple Sequence Comparison by Log-Expectation online tool to determine the 5’ and 3’ of exons used in construct design.

To design a construct containing two mutually exclusive exons, flanking introns of both exons were included in addition to an exon before the first mutually exclusive exon and an exon following the second mutually exclusive exon. This was done to avoid disturbing any spliceosome interactions with exons of interest and their corresponding 5’ and 3’ consensus splice sites. A reading frame of the first exon of a construct was determined by aligning an isoform’s cDNA sequence with the *cac* DNA, translating the resulting exon sequence into a peptide using Expasy online tool, and using the pBLAST program to confirm the resulting peptide sequence.

The next step was to introduce one- or two-base pair addition mutations within either of mutually exclusive exons avoiding proximity to splice sites. Introducing extra-base pairs within a spliced exon would result in a reading frame shift downstream of that exon. An introduction of extra-base pairs within a spliced exon may also result in a new stop codon, and so we avoided this by introducing several point mutations within the 3’ flanking exon of these constructs. Next, we used a 2A peptide sequence from porcine teschovirus-1 polyprotein between the 3’ exon and the first fluorophore and between the first and the second fluorophore. A GSG motif was added prior to each P2A sequence for improved cleavage of the resulting peptides. We added two base pairs after the first fluorophore to bring the second fluorophore into a reading frame.

An identical design strategy was employed for the constructs containing exons 5, and 6 as well as exons 10 and 11. A similar construct design was employed for exon 34 but without a corresponding mutually exclusive partner exon; exon 34 was flanked by 5’ and 3’ intronic material and exons 33 and 35. The original construct containing exon 34 had two one-base-pair missense mutations including A to T substitution at the 5’ +6 position of exon 35. From a *Drosophila* splice site predictor, G or T are highly unlikely to appear at this position, but to check this we also designed a construct identical to the original but with A to C point mutation to determine if the same fluorescent pattern would be observed with either construct. The data from the two different constructs were indistinguishable and only data from the former construct was included in the manuscript. None of the other constructs were re-synthetized due to their corresponding point mutations being at a significant distance from 5’ or 3’ of exons.

The resulting DNA constructs were flanked at the 5’ and 3’ by the restriction endonuclease sites NotI and AgeI respectively and cloned into the multiple cloning site downstream of the hsp70 promoter in the pJFRC14 plasmid.

### Construct Sequences in the order they appear in the manuscript

**cac_10_11_GFP_TagRFP-T** (Fig. 1F)

**cac_10_11_GFP_mRFP1** (Fig. 4B)

**cac_10_11_mRFP1_GFP** (Fig. 4F)

**cac_5_6_GFP_mRFP1** (Fig. 5C)

**cac_34_GFP_mRFP1** (Fig. 5H)

**cac_34_GFP_mRFP1_remade**

Nucleotide Sequences are available in a Supplemental File.

### Preparation of fillet-dissected larval for Confocal Imaging of Exon Reporters

All experiments were performed on female 3^rd^ instar larvae. The larvae were fillet dissected in cold Schneider’s insect medium on a Sylgard bath/tablet with an elevation at its center, elevating the middle of the dissection. The preparation was washed thrice with cold HL3 (0.1mM Ca^2+^, 15mM Mg^2+^), covered with a glass coverslip, and imaged with a Nikon 60X, 1.20 NA, Plan Apochromat VC water-immersion objective on a Nikon A1R confocal microscope fitted with GaAsP detectors. Preparations were scanned sequentially, starting with the longest wavelengths and progressing to the shortest (560 nm, 488 nm, 405 nm). All images were taken using the same settings. All images represent a collapsed Z-series encompassing a limited depth of the ventral ganglion or the full depth of terminal boutons in the periphery (3 μm and 1 μm step sizes, respectively).

### Confocal Imaging of Exon Reporters in Adult Brains

We selected 1-week-old adult female flies. They were dissected in cold Schneider’s insect medium, where the brain, along with the ventral nerve cord attached, was carefully extracted. The preparation was then transferred to the same Sylgard bath/tablet described above with the most anterior end “propped up” on the central elevation. A pin was placed over the cervical connective to hold it in place. The preparation was covered in a glass coverslip and imaged using the same settings as in larval preparations.

### Exon Reporter Imaging Data Analysis

We analyzed the images using ImageJ software. Measurements were obtained by calculating the average intensity from each collapsed z-series of images. ROIs were selected using a square of 5 × 5 pixels centrally positioned on a terminal bouton, glial process, muscle fiber, neuronal cell body or MB lobe. Care was taken to avoid the aggregates seen when using the mRFP1 fluorophore. Using the equation *VIII*, we can quantify the molar ratio of two exons by simply multiplying the correction coefficient χ to the ratio of corresponding fluorophores. Outliers were excluded if they fell outside limits defined by 2.5X the Median Absolute Deviation.

### Electron Microscopy

The number of AZs (N_AZ_) at each terminal was determined through a combination of light microscopy estimates of terminal volume (Justs et al., 2022), and transmission electron microscopy (TEM) estimates of the number of AZs per unit terminal volume. Five TEM series (100-nm sections), previously examined (Justs et al., 2022; Lu et al., 2016), were re-analyzed for the number of AZ profiles per unit volume. The type-Is terminals could be distinguished from type-Ib without ambiguity through reference to synaptic vesicle (SV) outer diameter (type-Is: 43.97±0.05nm; type-Ib: 33.67±0.05nm; mean±SEM), and each type-Ib MN could be distinguished from another without ambiguity through reference to muscle fiber identity. The minimum terminal volume sampled in each series was >1μm^3^ for type-Is and >5μm^3^ for type-Ib. Finally, N_AZ_ was determining by multiplying average terminal volume by the average number of AZs per unit volume (Table 1).

### Electrophysiology

Two-electrode voltage clamp (TEVC) was used to quantify terminal-specific release on each of body-wall muscle fibers #6, #13 and #12. Electrophysiology was conducted on female 3^rd^ instar larvae, from an in-house *w*^1118^ wildtype stock, in hemolymph-like solution #6 (HL6) (Macleod et al., 2002) containing MgCl_2_ added to 15 mM and CaCl_2_ added to 2 mM. Fillet dissections were performed in chilled HL6 on Sylgard plates and recordings were made 20 to 60 minutes after transecting the segmental nerves. Signals were detected, digitized and recorded using an Axoclamp 900A amplifier (Molecular Devices; Sunnyvale, CA) connected to a 4/35 PowerLab (ADInstruments; Colorado Springs, CO) and a PC running LabChart v8.0. Micropipettes filled with a 1:1 mixture of 3 M KCl and 3 M K-acetate. Measurements were performed on segment #4 under a 20X water-dipping objective of a BX50WI Olympus microscope to allow unequivocal identification of muscle fibers. Recordings commenced in current clamp mode, with two different micropipettes in two different muscle fibers. A suction pipette applied 0.3 ms electrical impulses to the transected nerve to evoke release from MN terminals. Impulse voltage was incrementally increased to initiate action potentials in one MN but not the other, and knowledge of the stereotypical innervation was relied upon to determine the identity of the MN terminal responsible for evoked release (Lu et al., 2016). Once stimulus thresholds were established one micropipette was removed and replaced in the same muscle fiber as the other micropipette and TEVC was initiated. A minimum of 10 Excitatory Junctional Currents (EJCs) were recorded during 0.2 Hz stimulation, along with 30 miniature EJCs (mEJCs), from each muscle fiber. Recordings of EJCs were made from 15 MN6-Ib terminals, 4 MN13-Ib, 24 MN12-Ib, 20 MNSNb/d-Is M#6, 9 MNSNb/d-Is M#13, 6 MNSNb/d-Is M#6, and mEJCs were recorded from 58 #6, 16 #13 and 61 #12 muscle fibers.

Quantal Content (QC) was calculated by dividing the mean EJC amplitude by the corrected mean mEJC amplitude. A correction factor must be applied to the mean mEJC amplitude to obtain a better estimate of terminal-specific QC. The correction is needed as the identity of the MN responsible for mEJCs is inscrutable using TEVC, yet mEJCs originating from type-Is terminals are 50% larger than those originating from type-Ib terminals (Dawson-Scully et al., 2007; Han et al., 2022; Karunanithi et al., 2002; Pawlu et al., 2004). As each terminal is responsible for a similar number of spontaneous events, a correction factor of 0.8 is applied to the mean mEJC amplitude (i.e. reduced by 20%) when calculating QC for type-Ib terminals, and a correction factor of 1.2 used for type-Is terminals (i.e. increased by 20%) (Lu et al., 2016).

Motor neuron endogenous firing frequencies were determined in 3^rd^ instar larvae during fictive locomotion as described by Chouhan and others (Chouhan et al., 2012; Chouhan et al., 2010). The firing frequencies for MN13-Ib and MNSNb/d-Is M#13 were determined by Chouhan et al., (2010), while those of MN6-Ib, MN13-Ib, MN12-Ib were determined by Chouhan et al., 2012.

### Ca^2+^ imaging and estimation of Ca^2+^ entry

Ca^2+^ imaging was performed as described by Justs and others (Justs et al., 2022) after forward filling MN terminals with a dextran-conjugated Ca^2+^ indicator (rhod) mixed with a dextran-conjugated Ca^2+^ insensitive dye (AF647). Data were collected on EMCCD cameras running at 100 frames-per-second. Justs and others reported the number of calcium ions (Ca^2+^) that enter each of the terminals with each action potential [Extended Data Set – Table 3-1 (Justs et al., 2022)]. In this study, we analyzed our electron microscopy data to estimate the number of AZs per unit terminal volume, and then used the terminal volume data from Justs and others to calculate the total number of AZs in each terminal. In the final step we divided the number of Ca^2+^ entering each terminal by the total number of AZs in each terminal, to calculate the number of Ca^2+^ entering each terminal per AZ per action potential.

### Statistical analysis and data presentation

Statistical tests were performed using SigmaStat 3.5 (integrated with SigmaPlot 10). Significance was assessed with an α of <0.05. Student’s t-tests were used for comparisons between two populations and Mann Whitney Rank Sum tests were used as a nonparametric alternative. α was adjusted according using the Bonferroni correction if more than one test was applied. Where Analysis of Variance (ANOVA) was used for multiple comparisons an overall α of <0.05 was required to claim significance (Fig 3). ANOVAs were run on ranks when tests for data normalcy failed. Propagation of uncertainty theory (Farrance & Frenkel, 2012) was used to calculate variance of means based on uncertainty measurements combined from different techniques. Pearson product-moment correlation coefficient was calculated to test the strength and direction of associations. The ordinary least-squares method was used to provide linear fits.

## Acknowledgements

This work was supported by NIH NINDS awards NS103906 and NS123377 to GTM. We are grateful for discussions with Drs Gil dos Santos, Scott Gratz, Karen Kim-Guisbert, Dushyant Mishra, Kate O’Connor-Giles and Mihaela Serpe.

