## Supplemental File for "Bichromatic exon-reporters reveal voltage-gated Ca^2+^-channel splice-isoform diversity across *Drosophila* neurons *in vivo*"

**Cac_10_11_GFP_TagRFP-T**

**GCGGCCGC**caaa**ATGtg**GACCAACGACGCATTAGGTTCAGCATTTAATTGGATATATTTCGTGCCTCTTA
TAGTTATAGGCTCATTTTTTATGCTCAACTTAGTTCTTGGTGTCCTTAGTGGGTAAGTTGATCGCGAATTGCT
TTATTTTAACGATGAAGCCACTTCTACACCAGTTCACATTATGTTCTACTTATAATTTCGATTACTTCCGTTGA
GAAAGAGAGTAAAAAAAACAAAAAACAAAAAAAAAAAAAAAATATGGAAAAATGGGATCTGAAAATACTG
CATACATATGTTTAGATCATTTCTATATGGAATGTCGTCGTGTGTTTTTTATGACGCATTTAAATATTTTTGTAA
AGATGAAATGTAAAGAAAGTGTGTCCAATTGCCAGATTATACATATGTAATATATCGGTTTTACGTAGAGCTA
CGTGCATGTGCATACACGGTGAAAAAAAAAAATGATAAAAAAATAAGTCAATTTGGCTGTAAATTATTCAAAAT
TTTCGCCTGTTGCCGGACGATTGAAATATATGTATGTACGTGTTTGATATACTGTCATTTTGATTTGCATTCAA
CTTGTACGTTTCGTTATATATTTGTTTTTATCTTTTTCCGAAATTTTGTTCAATGCATTTAACTATCGTATTCCCG
AATTCA**GA**GAGTTTTCGAACGAACGAAATCGTGTCGAACGTCGCATGGAGTTTCAAAAATGCCGCTTTCG
GGCCATGTTTCAGACAGCAATGGTCTCG**TA**CCTTcGACTGGATCACACAAGCAGGTTTATACAACTAGATCT
ATATCTAAGTGCAACCAAGTAAACTGCCATAATCAACCCGAACGTCATACAACACAACCTGCAATTATAACTA
ATTATAACTATTATCGGCCAAAAAAAAACAGAACAATAAGTCAACTCGTATCGTAATGCTAATGCTAATGCTA
TCGAATCCGAGCGAGTTCATTGAGCTAAACACCCATCTAATTAAACCAAAAAAAAAAACAATTCCGACCTAT
ACTTCCTTAACCAGTTTATAAAGAACGTGTTTGATTTACTGCTAGTTATGTATTCTTAAATAGATCGCAAGCA
ATTCCCCGGCTACAAAGAGTTCTGGTACCCAAAACTAACTTAACTATCAATTACAATACAACTACTATGTACT
TCCAATGAACCGAATTAATAATATTAACTGCGAAATGCCGTTCACCCCACTATAGTTTCTAACTAACCAACCA
ACCAAACAATCAATAATCTAATCCATATATACTCTGCAGTGCGTAATGCATATTATATTCTCTGCATTTTGCCAC
ACACTTAACCCGATTTTTACTACCATAGCGATAAAGTTGAATTCCCCCCCCCTCCCTGTCAAAATAAACCAAA
CTAACCAAATACATAGCTATTTCAACTGCAGTTAGCCAAATTAACCTCCAGTTATATACTTATACCCGCCAGCG
GACTCGTGAATTGTTGTTGAGTCTCGGCCTCGTTCTCGATATCTTGGCTAATATAGTACACTCCCTATACACTA
TATACACTGATAGAACAGTGCACCCACCTATCAATATCAATGTTCTGTCCCCGGTTAACTATCGCCATCAAAAT
GTATCATCGCTCCAGACTCAGAGACTTCAAGACTCGCAGACTCCCAGCGAATCAGGGCAAAATTCAGAGT
TATTCAAGACCAGATGACGCTATGGCATCATCACCCAATGCGTACTCTATATATTTATTTGTCGCCACCGAATA
AATCATTTTATATATGCTTGTATGCACACATATAATCTAACACATTTCCCTTCTTTTATATATATTTTTCCTTGACT
TTTGACCTCATTGAAGGTTCCCGTCGAGGAGGCGCAATAAATATATACATATATATACAATATATGTGTTAAAT
TGCATTTCTGTTACTTCATTTCCAGTAAATGATTTCGACTTTCTGTTTATGCCCGTTTGTTGTGTTCTAAAAAA
CAAAAAAAAAAAACAAAGCACTGGAATATTCTTGATTTTACGAATCTTTAAGTTATTCTTCTCTGTTGATTACA
CTCGTACAAAAACAATGGCGAAAGTATGATGTGAACTTTGATATTGTAATACTGAATACCAAAGCATACATG
ATCGTTTTAGCAATCAATCAATCCTGTTTTTTTGGTTTATCACGTTCTATTTAATTATTTTTTGTA**GT**GAATTCG
C**AAAAGAACGAGAAA**AAGTAGAAAATAGACAAGAGTTTCTTAAACTTAGAAGGCAGCAGCAACTAGAA
AGAGAGTTAAACGGCTATGTTGAATGGATTTGTAAAG**CTG**GTATAGTATTTACCCCTATTTCTTGTTTGTAAA
TCTATATATACACACACACACAATGACTAAACTCATGAATAACTCTCATACACACATATATACAAACATAAAAA
TTTAAACACTATGCGCCCCCATATGTTCATCCACATCCATATGTACATATATATATAAATTATAAAAACGACATTT
ATATCAGGTCCTTTTCTATGACTATAAAACTGAATTCAATGAAAATAGGCCAACTATAATCACATAAAATATTT
TTTTAAAAATTCGAACAGGGATCACAAAAAATGCCTAACAAAAGATTCTTCTTTAATGTCATTGCTTCACTG
AAAGATACATTAACTAAATATCGTTTATATCATTTTCATATAAAATGACGATATATTCATGCATATTTAAAGTGAT
ACAAGAATAAAATATACGTTGTATTGAGATGTTCCCTAATAGCCAGCACACTTTTGGCACATTCTTAGCATCC
CTCTCCTCATAGAATTGAAACCCTCGAACCTATATACATTGCCAAATTTGAAAGTGAATAAATTCCTCAATATA
CTCAGAATTAAAATAGTTTTCGTGGAAATATGGACATATATCCCTCATGCCGCCCCCAATATATATACATCTAC
CCACATTGACTATAATGTACATACGAGTATATATCCTTTCTGCTTATGTGTTGATCAAATCGATTCCCTTATTGT
GACCCATCACCAATCACTCGATTTTAATGCTATTTCAGAGGAGGTAATCCTGGCCGAGGAGCGCACCACGG

AAGAAGAGAAAATGCACATAATGGAgGGAAGCGGAgccaccaacttctccctgctgaagcaggccggcgacgtggagga
gaaccccggccccatggtgagcaagggcgaggagctgttcaccggggtggtgcccatcctggtcgagctggacggcgacgtaaacggcc
acaagttcagcgtgtccggcgagggcgagggcgatgccacctacggcaagctgaccctgaagttcatctgcaccaccggcaagctgcccg
tgccctggcccaccctcgtgaccaccctgacctacggcgtgcagtgcttcagccgctaccccgaccacatgaagcagcacgacttcttcaa
gtccgccatgcccgaaggctacgtccaggagcgcaccatcttcttcaaggacgacggcaactacaagacccgcgccgaggtgaagttcga
gggcgacaccctggtgaaccgcatcgagctgaagggcatcgacttcaaggaggacggcaacatcctggggcacaagctggagtacaac
tacaacagccacaacgtctatatcatggccgacaagcagaagaacggcatcaaggtgaacttcaagatccgccacaacatcgaggacgg
cagcgtgcagctcgccgaccactaccagcagaacacccccatcggcgacggccccgtgctgctgcccgacaaccactacctgagcaccc
agtccgccctgagcaaagaccccaacgagaagcgcgatcacatggtcctgctggagttcgtgaccgccgccgggatcactctcggcatgg
acgagctgtacaag**taa**AAGGAAGCGGAgccaccaacttctccctgctgaagcaggccggcgacgtggaggagaaccccggcccc
gtgtctaagggcgaagagctgattaaggagaacatgcacatgaagctgtacatggagggcaccgtgaacaaccacca
cttcaagtgcacatccgagggcgaaggcaagccctacgagggcacccagaccatgagaatcaaggtggtcgagggc
ggccctctcccctcgccttcgacatcctggctaccagcttcatgtacggcagcagaaccttcatcaaccacacccagggc
atccccgatttctttaagcagtccttccctgagggcttcacatgggagagagtcaccacatacgaagacgggggcgtgctg
accgctacccaggacaccagcctccaggacggctgcctcatctacaacgtcaagatcagaggggtgaacttcccatcca
acggccctgtgatgcagaagaaaacactcggctgggaggccaacaccgagatgctgtaccccgctgacggcggcctg
gaaggcagaaccgacatggccctgaagctcgtgggcgggggccacctgatctgcaacttcaagaccacatacagatcc
aagaaacccgctaagaacctcaagatgcccggcgtctactatgtggaccacagactggaaagaatcaaggaggccga
caaagagacctacgtcgagcagcacgaggtggctgtggccagatactgcgacctccctagcaaactggggcacaaac
ttaatggcatggacgagctgtacaagtaa**ACCGGT**

**Cac_10_11_GFP_mRFP1**

**GCGGCCGC**caaa**ATG**tgGACCAACGACGCATTAGGTTCAGCATTTAATTGGATATAT
TTCGTGCCTCTTATAGTTATAGGCTCATTTTTTATGCTCAACTTAGTTCTTGGTGTCC
TTAGTGGGTAAGTTGATCGCGAATTGCTTTATTTTAACGATGAAGCCACTTCTACAC
CAGTTCACATTATGTTCTACTTATAATTTCGATTACTTCCGTTGAGAAAGAGAGTAA
AAAAAACAAAAAACAAAAAAAAAAAAAAAATATGGAAAAATGGGATCTGAAAATACTG
CATACATATGTTTAGATCATTTCTATATGGAATGTCGTCGTGTGTTTTTTATGACGCA
TTTAAATATTTTTGTAAAGATGAAATGTAAAGAAAGTGTGTCCAATTGCCAGATTATA
CATATGTAATATATCGGTTTTACGTAGAGCTACGTGCATGTGCATACACGGTGAAA
AAAAAAATGATAAAAAATAAGTCAATTTGGCTGTAAATTATTCAAAATTTTCGCCTGT
TGCCGGACGATTGAAATATATGTATGTACGTGTTTGATATACTGTCATTTTGATTTG
CATTCAACTTGTACGTTTCGTTATATATTTGTTTTTATCTTTTTCCGAAATTTTGTTCA
ATGCATTTAACTATCGTATTCCCGAATTCA**GA**GAGTTTTCGAACGAACGAAATCGTG
TCGAACGTCGCATGGAGTTTCAAAAATGCCGCTTTCGGGCCATGTTTCAGACAGCA
ATGGTCTCG**TA**CCTTcGACTGGATCACACAAGCAGGTTTATACAACTAGATCTATAT
CTAAGTGCAACCAAGTAAACTGCCATAATCAACCCGAACGTCATACAACACAACCT
GCAATTATAACTAATTATAACTATTATCGGCCAAAAAAAAACAGAACAATAAGTCAA
CTCGTATCGTAATGCTAATGCTAATGCTATCGAATCCGAGCGAGTTCATTGAGCTA
AACACCCATCTAATTAAACCAAAAAAAAAAAACAATTCCGACCTATACTTCCTTAACC
AGTTTATAAAGAACGTGTTTGATTTACTGCTAGTTATGTATTCTTAAATAGATCGCAA
GCAATTCCCCGGCTACAAAGAGTTCTGGTACCCAAAACTAACTTAACTATCAATTAC
AATACAACTACTATGTACTTCCAATGAACCGAATTAATAATATTAACTGCGAAATGC

CGTTCACCCCACTATAGTTTCTAACTAACCAACCAACCAAACAATCAATAATCTAAT
CCATATATACTCTGCAGTGCGTAATGCATATTATATTCTCTGCATTTTGCCACACAC
TTAACCCGATTTTTACTACCATAGCGATAAAGTTGAATTCCCCCCCCCTCCCTGTCA
AAATAAACCAAACTAACCAAATACATAGCTATTTCAACTGCAGTTAGCCAAATTAAC
CTCCAGTTATATACTTATACCCGCCAGCGGACTCGTGAATTGTTGTTGAGTCTCGG
CCTCGTTCTCGATATCTTGGCTAATATAGTACACTCCCTATACACTATATACACTGA
TAGAACAGTGCACCCACCTATCAATATCAATGTTCTGTCCCCGGTTAACTATCGCC
ATCAAAATGTATCATCGCTCCAGACTCAGAGACTTCAAGACTCGCAGACTCCCAGC
GAATCAGGGCAAAATTCAGAGTTATTCAAGACCAGATGACGCTATGGCATCATCAC
CCAATGCGTACTCTATATATTTATTTGTCGCCACCGAATAAATCATTTTATATATGCT
TGTATGCACACATATAATCTAACACATTTCCCTTCTTTTATATATATTTTTCCTTGACT
TTTGACCTCATTGAAGGTTCCCGTCGAGGAGGCGCAATAAATATATACATATATATA
CAATATATGTGTTAAATTGCATTTCTGTTACTTCATTTCCAGTAAATGATTTCGACTT
TCTGTTTATGCCCGTTTGTTGTGTTCTAAAAAACAAAAAAAAAAAACAAAGCACTGGA
ATATTCTTGATTTTACGAATCTTTAAGTTATTCTTCTCTGTTGATTACACTCGTACAA
AAACAATGGCGAAAGTATGATGTGAACTTTGATATTGTAATACTGAATACCAAAGCA
TACATGATCGTTTTAGCAATCAATCAATCCTGTTTTTTTGGTTTATCACGTTCTATTT
AATTATTTTTTGTA**GT**GAATTCGC**AAAAGAACGAGAAA**AAGTAGAAAATAGACAAG
AGTTTCTTAAACTTAGAAGGCAGCAGCAACTAGAAAGAGAGTTAAACGGCTATGTT
GAATGGATTTGTAAAG**CTG**GTATAGTATTTACCCCTATTTCTTGTTTGTAAATCTATA
TATACACACACACACAATGACTAAACTCATGAATAACTCTCATACACACATATATAC
AAACATAAAAATTTAAACACTATGCGCCCCCATATGTTCATCCACATCCATATGTAC
ATATATATATAAATTATAAAAACGACATTTATATCAGGTCCTTTTCTATGACTATAAAA
CTGAATTCAATGAAAATAGGCCAACTATAATCACATAAAATATTTTTTTAAAAATTCG
AACAGGGATCACAAAAAATGCCTAACAAAAGATTCTTCTTTAATGTCATTGCTTCAC
TGAAAGATACATTAACTAAATATCGTTTATATCATTTTCATATAAAATGACGATATATT
CATGCATATTTAAAGTGATACAAGAATAAAATATACGTTGTATTGAGATGTTCCCTA
ATAGCCAGCACACTTTTGGCACATTCTTAGCATCCCTCTCCTCATAGAATTGAAACC
CTCGAACCTATATACATTGCCAAATTTGAAAGTGAATAAATTCCTCAATATACTCAG
AATTAAAATAGTTTTCGTGGAAATATGGACATATATCCCTCATGCCGCCCCCAATAT
ATATACATCTACCCACATTGACTATAATGTACATACGAGTATATATCCTTTCTGCTTA
TGTGTTGATCAAATCGATTCCCTTATTGTGACCCATCACCAATCACTCGATTTTAAT
GCTATTTCAGAGGAGGTAATCCTGGCCGAGGAGCGCACCACGGAAGAAGAGAAAA
TGCACATAATGGAgGGAAGCGGAgccaccaacttctccctgctgaagcaggccggcgacgtggaggag
aaccccggccccatggtgagcaagggcgaggagctgttcaccggggtggtgcccatcctggtcgagctggacggcgac
gtaaacggccacaagttcagcgtgtccggcgagggcgagggcgatgccacctacggcaagctgaccctgaagttcatc
tgcaccaccggcaagctgcccgtgccctggcccaccctcgtgaccaccctgacctacggcgtgcagtgcttcagccgcta
ccccgaccacatgaagcagcacgacttcttcaagtccgccatgcccgaaggctacgtccaggagcgcaccatcttcttca
aggacgacggcaactacaagacccgcgccgaggtgaagttcgagggcgacaccctggtgaaccgcatcgagctgaa
gggcatcgacttcaaggaggacggcaacatcctggggcacaagctggagtacaactacaacagccacaacgtctatat
catggccgacaagcagaagaacggcatcaaggtgaacttcaagatccgccacaacatcgaggacggcagcgtgcag
ctcgccgaccactaccagcagaacacccccatcggcgacggccccgtgctgctgcccgacaaccactacctgagcac
ccagtccgccctgagcaaagaccccaacgagaagcgcgatcacatggtcctgctggagttcgtgaccgccgccgggat
cactctcggcatggacgagctgtacaag**taa**AAGGAAGCGGAgccaccaacttctccctgctgaagcaggccgg
cgacgtggaggagaaccccggccccatggcctcctccgaggacgtcatcaaggagttcatgcgcttcaaggtgcgcatg
gagggctccgtgaacggccacgagttcgagatcgagggcgagggcgagggccgcccctacgagggcacccagacc

gccaagctgaaggtgaccaagggcggcccctgcccttcgcctgggacatcctgtcccctcagttccagtacggctccaa
ggcctacgtgaagcaccccgccgacatccccgactacttgaagctgtccttccccgagggcttcaagtgggagcgcgtg
atgaacttcgaggacggcggcgtggtgaccgtgacccaggactcctccctgcaggacggcgagttcatctacaaggtga
agctgcgcggcaccaacttcccctccgacggccccgtaatgcagaagaagaccatgggctgggaggcctccaccgag
cggatgtaccccgaggacggcgccctgaagggcgagatcaagatgaggctgaagctgaaggacggcggccactacg
acgccgaggtcaagaccacctacatggccaagaagcccgtgcagctgcccggcgcctacaagaccgacatcaagct
ggacatcacctcccacaacgaggactacaccatcgtggaacagtacgagcgcgccgagggccgccactccaccggc
gcctaa**ACCGGT**

**Cac_10_11_mRFP1_GFP**

**GCGGCCGC**caaa**ATG**tgGACCAACGACGCATTAGGTTCAGCATTTAATTGGATATAT
TTCGTGCCTCTTATAGTTATAGGCTCATTTTTTATGCTCAACTTAGTTCTTGGTGTCC
TTAGTGGGTAAGTTGATCGCGAATTGCTTTATTTTAACGATGAAGCCACTTCTACAC
CAGTTCACATTATGTTCTACTTATAATTTCGATTACTTCCGTTGAGAAAGAGAGTAA
AAAAAACAAAAAACAAAAAAAAAAAAAAAATATGGAAAAATGGGATCTGAAAATACTG
CATACATATGTTTAGATCATTTCTATATGGAATGTCGTCGTGTGTTTTTTATGACGCA
TTTAAATATTTTTGTAAAGATGAAATGTAAAGAAAGTGTGTCCAATTGCCAGATTATA
CATATGTAATATATCGGTTTTACGTAGAGCTACGTGCATGTGCATACACGGTGAAA
AAAAAAATGATAAAAAATAAGTCAATTTGGCTGTAAATTATTCAAAATTTTCGCCTGT
TGCCGGACGATTGAAATATATGTATGTACGTGTTTGATATACTGTCATTTTGATTTG
CATTCAACTTGTACGTTTCGTTATATATTTGTTTTTATCTTTTTCCGAAATTTTGTTCA
ATGCATTTAACTATCGTATTCCCGAATTCA**GA**GAGTTTTCGAACGAACGAAATCGTG
TCGAACGTCGCATGGAGTTTCAAAAATGCCGCTTTCGGGCCATGTTTCAGACAGCA
ATGGTCTCG**TA**CCTTcGACTGGATCACACAAGCAGGTTTATACAACTAGATCTATAT
CTAAGTGCAACCAAGTAAACTGCCATAATCAACCCGAACGTCATACAACACAACCT
GCAATTATAACTAATTATAACTATTATCGGCCAAAAAAAAACAGAACAATAAGTCAA
CTCGTATCGTAATGCTAATGCTAATGCTATCGAATCCGAGCGAGTTCATTGAGCTA
AACACCCATCTAATTAAACCAAAAAAAAAAAACAATTCCGACCTATACTTCCTTAACC
AGTTTATAAAGAACGTGTTTGATTTACTGCTAGTTATGTATTCTTAAATAGATCGCAA
GCAATTCCCCGGCTACAAAGAGTTCTGGTACCCAAAACTAACTTAACTATCAATTAC
AATACAACTACTATGTACTTCCAATGAACCGAATTAATAATATTAACTGCGAAATGC
CGTTCACCCCACTATAGTTTCTAACTAACCAACCAACCAAACAATCAATAATCTAAT
CCATATATACTCTGCAGTGCGTAATGCATATTATATTCTCTGCATTTTGCCACACAC
TTAACCCGATTTTTACTACCATAGCGATAAAGTTGAATTCCCCCCCCCTCCCTGTCA
AAATAAACCAAACTAACCAAATACATAGCTATTTCAACTGCAGTTAGCCAAATTAAC
CTCCAGTTATATACTTATACCCGCCAGCGGACTCGTGAATTGTTGTTGAGTCTCGG
CCTCGTTCTCGATATCTTGGCTAATATAGTACACTCCCTATACACTATATACACTGA
TAGAACAGTGCACCCACCTATCAATATCAATGTTCTGTCCCCGGTTAACTATCGCC
ATCAAAATGTATCATCGCTCCAGACTCAGAGACTTCAAGACTCGCAGACTCCCAGC
GAATCAGGGCAAAATTCAGAGTTATTCAAGACCAGATGACGCTATGGCATCATCAC
CCAATGCGTACTCTATATATTTATTTGTCGCCACCGAATAAATCATTTTATATATGCT
TGTATGCACACATATAATCTAACACATTTCCCTTCTTTTATATATATTTTTCCTTGACT
TTTGACCTCATTGAAGGTTCCCGTCGAGGAGGCGCAATAAATATATACATATATATA

CAATATATGTGTTAAATTGCATTTCTGTTACTTCATTTCCAGTAAATGATTTCGACTT
TCTGTTTATGCCCGTTTGTTGTGTTCTAAAAAACAAAAAAAAAAAACAAAGCACTGGA
ATATTCTTGATTTTACGAATCTTTAAGTTATTCTTCTCTGTTGATTACACTCGTACAA
AAACAATGGCGAAAGTATGATGTGAACTTTGATATTGTAATACTGAATACCAAAGCA
TACATGATCGTTTTAGCAATCAATCAATCCTGTTTTTTTGGTTTATCACGTTCTATTT
AATTATTTTTTGTA**GT**GAATTCGC**AAAAGAACGAGAAA**AAGTAGAAAATAGACAAG
AGTTTCTTAAACTTAGAAGGCAGCAGCAACTAGAAAGAGAGTTAAACGGCTATGTT
GAATGGATTTGTAAAG**CTG**GTATAGTATTTACCCCTATTTCTTGTTTGTAAATCTATA
TATACACACACACACAATGACTAAACTCATGAATAACTCTCATACACACATATATAC
AAACATAAAAATTTAAACACTATGCGCCCCCATATGTTCATCCACATCCATATGTAC
ATATATATATAAATTATAAAAACGACATTTATATCAGGTCCTTTTCTATGACTATAAAA
CTGAATTCAATGAAAATAGGCCAACTATAATCACATAAAATATTTTTTTAAAAATTCG
AACAGGGATCACAAAAAATGCCTAACAAAAGATTCTTCTTTAATGTCATTGCTTCAC
TGAAAGATACATTAACTAAATATCGTTTATATCATTTTCATATAAAATGACGATATATT
CATGCATATTTAAAGTGATACAAGAATAAAATATACGTTGTATTGAGATGTTCCCTA
ATAGCCAGCACACTTTTGGCACATTCTTAGCATCCCTCTCCTCATAGAATTGAAACC
CTCGAACCTATATACATTGCCAAATTTGAAAGTGAATAAATTCCTCAATATACTCAG
AATTAAAATAGTTTTCGTGGAAATATGGACATATATCCCTCATGCCGCCCCCAATAT
ATATACATCTACCCACATTGACTATAATGTACATACGAGTATATATCCTTTCTGCTTA
TGTGTTGATCAAATCGATTCCCTTATTGTGACCCATCACCAATCACTCGATTTTAAT
GCTATTTCAGAGGAGGTAATCCTGGCCGAGGAGCGCACCACGGAAGAAGAGAAAA
TGCACATAATGGAgGGAAGCGGAgccaccaacttctccctgctgaagcaggccggcgacgtggaggag
aaccccggccccatggcctcctccgaggacgtcatcaaggagttcatgcgcttcaaggtgcgcatggagggctccgtga
acggccacgagttcgagatcgagggcgagggcgagggccgcccctacgagggcacccagaccgccaagctgaagg
tgaccaagggcggccccctgcccttcgcctgggacatcctgtcccctcagttccagtacggctccaaggcctacgtgaag
caccccgccgacatccccgactacttgaagctgtccttccccgagggcttcaagtgggagcgcgtgatgaacttcgagga
cggcggcgtggtgaccgtgacccaggactcctccctgcaggacggcgagttcatctacaaggtgaagctgcgcggcac
caacttcccctccgacggccccgtaatgcagaagaagaccatgggctgggaggcctccaccgagcggatgtaccccg
aggacggcgccctgaagggcgagatcaagatgaggctgaagctgaaggacggcggccactacgacgccgaggtca
agaccacctacatggccaagaagcccgtgcagctgcccggcgcctacaagaccgacatcaagctggacatcacctcc
cacaacgaggactacaccatcgtggaacagtacgagcgcgccgagggccgccactccaccggcgcctaaAAGGA
AGCGGAgccaccaacttctccctgctgaagcaggccggcgacgtggaggagaaccccggccccAtggtgagcaa
gggcgaggagctgttcaccggggtggtgcccatcctggtcgagctggacggcgacgtaaacggccacaagttcagcgt
gtccggcgagggcgagggcgatgccacctacggcaagctgaccctgaagttcatctgcaccaccggcaagctgcccgt
gccctggcccaccctcgtgaccaccctgacctacggcgtgcagtgcttcagccgctaccccgaccacatgaagcagcac
gacttcttcaagtccgccatgcccgaaggctacgtccaggagcgcaccatcttcttcaaggacgacggcaactacaaga
cccgcgccgaggtgaagttcgagggcgacaccctggtgaaccgcatcgagctgaagggcatcgacttcaaggaggac
ggcaacatcctggggcacaagctggagtacaactacaacagccacaacgtctatatcatggccgacaagcagaagaa
cggcatcaaggtgaacttcaagatccgccacaacatcgaggacggcagcgtgcagctcgccgaccactaccagcaga
acacccccatcggcgacggccccgtgctgctgcccgacaaccactacctgagcacccagtccgccctgagcaaagac
cccaacgagaagcgcgatcacatggtcctgctggagttcgtgaccgccgccgggatcactctcggcatggacgagctgt
acaagtaa**ACCGGT**

**Cac_5_6_GFP_mRFP1**

**<u>GCGGCCGC</u>**caaaATGttGGAAAAAACGGAGGCCTATTTTTATGCATTTTCTGTGTAG
AAGCGTCGCTCAAGATCCTCGCCTTAGGGCTTGTTCTGCATAAACACTCCTATCTC
AGGAATATTTGGAACATCATGGATTTTTTCGTTGTAGTTACGGGGTAAATAATGAAC
CAAACTATTTTCGAAACATATGCACCAATTTTATTTTCAACTAATATCGTATTAAAAA
AAAAACTACACGCCGTGCAAGCATTATCTGTTTTTATTCAAACCAAACCAATATCAC
CTATGTTTAAGTCTTGATCATAATGAATCCCAAGAAAAAGTTAACTTAATATTCTCAG
AAAAGCTTAGTTACTCTTTTTCTAGATTCATTACAGATCTAATCTAACAATTAGACAT
GTCATTTTAGTTGTAAGATGTAAAATTGTACAAATCACAAACTTTATATTAAGATATG
TATTTAGTTATTAAGTGCCCAATATATTTTTGTGTTGCGGTAAGATATTTGCCTCAAC
TAAATCATATAAATGTATTATAAGTAAGCGACTATATTAACAACTTCTCCCATATATG
TTTAAAATCGGGTAACAAACTCGAAATGGCAAATAACAATGCCAATTGCAGAGCCA
TGACGATATTTGCTGAGGCCAATATAGATGTTGACCTGCGTATGTTACGATCTTTTC
GTGTTTTACGCCCACTGAAGCTCGTATCCCGAATTCCAAGTAAGACATCTGCCAAC
AAAAAGAACCATTAGTTGCTACTACCAAAAACCTAGGCCAGATAAAAAAAACATCGT
TTTCTTCAGTAAACATATATATTAAACTAAACAAATTACACTCGAAAAAGCTAGTTGT
CCTATATGGATACTTACTATAAATTTAATTGGATTAGGAGTAGCGAAAACGGCATAA
ACAGAAAACAACCTTAGAGGCTAGGAGCCACGAAAATTAGATGCTAAAATATATTTA
TTAGTTATATATTAAGTAGCTACATATAATATAAATTTATAGCTCAATCAGGCTTCCG
ATTTCCTTTGATTCTTCAAATGAAACAAAATAAATTTTATTAAGGAAAACGGTAGCAC
GTTTGTAAAATTCTCTTATAAACAAAATAGGATTAGTATTATTCAACTATAGATGGCA
TCAGTAGGATGCAATGTGTTATCGTGTAGGTTTATAATATAAATCAATTCATGTACA
GTGGTAGCCACGAATTTAGGAACTTTTAAAGCGTATTATCTTTCGTGGCTATCACTC
GGTAGTTTTTATGAATATATATATATATACATATAGCTGTATATTCTTTCTGATTTGTA
GTGATTTCGACTAGCCATTTTTATTTATGTTTACAAAATATTATATTAGCATTATAATT
ACTAGATATATATATACATAAGTGAGTCAAAGACTATATAACTGTGTGTGATTCTCG
CGACTATAATGTTTATAAATTAGACGTTAAGTCTTGACCGTAATTCAAGTTACCTTTC
CGTTCGATTTAAAATAGATCGCAACTTGCAAGTCATCGATGAATCCCCTTTTAAAGG
TGGTAGAATAATAGCGAATATTATTGATTTTCGATAATAGCTGGCACATAAATGATA
AATTTACATCTATATATAATGCCGAAACTATAGAGTGACTGACCAAAATCACTTTTA
GCCAATTGTAGATGTTTTCATTTATTTTTCATCGGAAATTTGTATCTCTTAAAGTGAA
AATTTACTTTTACTCTCCTAGAACTTTGAAATAAGCAGTGTGTATGTTATAAGCAATA
GATTCATGTTATATTTTAAATTATAGCCTTAACCTATATTCAATTCCATAGCTGGACC
AAAACCATAGTTTACATTTGCTCTTCAACATTAATGCCTGAAAATATTGATCAATATT
GTTCGGAATGAAGAGCAAAGCTAAACTATTCTGCGTTTTCGCTTGGGTTACCCATT
GTGTGTGCGAAAAATATGAACTAACCGCGCGTTAAACGTTATTATTATTAAGTATTA
TTTAAGTTATTTAAATTTGTACACTCCTTAGTTGCTTCTTGCTTTCGATTCAAGTTTC
GATTTAAGCTGCACAATACATATGTATAAACTTTTGTTTTCAACGTACTCATATGATT
TCGATATTCCAATTATGATTATCCGATCTGTATCCATTGAGTATCGCTAATCAATATA
TAACTTCATTTCGATCGCTTAACGTCTGCAAACAGATTCATGACACAGTACCCACAA
ATAGGGCCCGAGGTAGACCTAAGAACACTTAGAGCCATTCGTGTGCTACGGCCCT
TAAAATTaaAGTGTCTGGAATTCCTAGTGAGTAGTTCCTCTGTTTAATTCTACATGTC
GTTGTTGTCTTTCCATTCTTGATTTTTAAAGCCAAACCTCACTCCAGACTCTTTAATG
TGCGTAAATGAGATTGTTTTTTTTTTTAAATTGTGTTTTAAGTTCTAGTACGAGTGAG
TGGAAAACGAGTGAGAAAAGAGCCTGCATTTTTGCCACCACTTAGTTTTCGATTTTC
GTTTGAAACGTACTCTTTTTGACTAGAATGATTGTTTGTGGCACATGATTGTCGTGG
AGGAGCTTTCAATGTCTGCGACAATCATGTCCTAAATGTGAAACACATACCCAGCA

TATGCATCTTTCTAATCCAACGTCCATCTTCTCCAAACAACTAACCAATCAACTTAA
CTGAATCGTTTCTGGATCGAATCGGAACGGCAACGTCAACAGGTTTACAAGTAcTT
TTAcAATCTATATTCAAGGCGATGGCACCTTTACTGCAAATCGGTCTCTTGGTGTTG
TTTGCAATCGTATTTTTTGCAATCATTGGACTCGAGTTTTATTCGGGCGCACTGCAT
AAGACTTGTTATAGCTCAGAAGATCCAAGGAAGCGGAgccaccaacttctccctgctgaagcag
gccggcgacgtggaggagaaccccggccccatggtgagcaagggcgaggagctgttcaccggggtggtgcccatcct
ggtcgagctggacggcgacgtaaacggccacaagttcagcgtgtccggcgagggcgagggcgatgccacctacggc
aagctgaccctgaagttcatctgcaccaccggcaagctgcccgtgccctggcccaccctcgtgaccaccctgacctacgg
cgtgcagtgcttcagccgctaccccgaccacatgaagcagcacgacttcttcaagtccgccatgcccgaaggctacgtcc
aggagcgcaccatcttcttcaaggacgacggcaactacaagacccgcgccgaggtgaagttcgagggcgacaccctg
gtgaaccgcatcgagctgaagggcatcgacttcaaggaggacggcaacatcctggggcacaagctggagtacaacta
caacagccacaacgtctatatcatggccgacaagcagaagaacggcatcaaggtgaacttcaagatccgccacaaca
tcgaggacggcagcgtgcagctcgccgaccactaccagcagaacacccccatcggcgacggccccgtgctgctgccc
gacaaccactacctgagcacccagtccgccctgagcaaagaccccaacgagaagcgcgatcacatggtcctgctgga
gttcgtgaccgccgccgggatcactctcggcatggacgagctgtacaagtaaAAGGAAGCGGAgccaccaactt
ctccctgctgaagcaggccggcgacgtggaggagaaccccggccccAtggcctcctccgaggacgtcatcaaggagtt
catgcgcttcaaggtgcgcatggagggctccgtgaacggccacgagttcgagatcgagggcgagggcgagggccgcc
cctacgagggcacccagaccgccaagctgaaggtgaccaagggcggccccctgcccttcgcctgggacatcctgtccc
ctcagttccagtacggctccaaggcctacgtgaagcaccccgccgacatccccgactacttgaagctgtccttccccgag
ggcttcaagtgggagcgcgtgatgaacttcgaggacggcggcgtggtgaccgtgacccaggactcctccctgcaggac
ggcgagttcatctacaaggtgaagctgcgcggcaccaacttcccctccgacggccccgtaatgcagaagaagaccatg
ggctgggaggcctccaccgagcggatgtaccccgaggacggcgccctgaagggcgagatcaagatgaggctgaagc
tgaaggacggcggccactacgacgccgaggtcaagaccacctacatggccaagaagcccgtgcagctgcccggcgc
ctacaagaccgacatcaagctggacatcacctcccacaacgaggactacaccatcgtggaacagtacgagcgcgccg
agggccgccactccaccggcgccTaa**<u>ACCGGT</u>**

**Cac_34_GFP_mRFP1**

**<u>GCGGCCGC</u>**caaaATGAGGCACAATGGCTCTCCACTGGCCAGATCTCCGAGTCCTC
GACGGCGTGGCCATCAATACATACATCATGATATCGGGTTCTCCGATACCGTATCT
AATGTTGTAGAGATGGTCAAGGAGACTCGTCATCCTAGGCATGGCAACAGTCATCC
GCGGTATCCAAGAGGTATTTATTGGCTCGACATCCATGGCATTTAATCATTAACTAC
TTTCAACTGACTATCGTGTGTGTGTGTGTGTGTGCTGGTTGTAAGCCAAATGATCT
GGTTAGGGATTCGTCAACATTATCCTGTTATCGAGTTGTCTGTCCTTGATGTTGCCC
TTAATCACTTCGTATCCATATGAGATCGGCTTGCAAGGGGCACACACAGAAAACAA
AGGGACTTGGACATTCCAGGGATTTTTGATCTATTTTGTAGAACCGTAGTTATATAC
ATATATACATATATGTCTACAATTCCATAGCTATTAACGAAATCTTTTAATACAAATAT
ATTATATTTCAATAATAATTTTATCGTTTCCGTTAAGTCGGTCCTCTATTGTCGTTTAT
CGTTTATCTCTTCGAAAGCAATTTGAAAATCGCAAAAAACCAAAATGAAAAAAAAAA
AAACAAAACAAAACAAAACAAAAGAAGAGAAGAGAAAATATGTTGCTAAGATCCAAT
ACGTAAAATGTTGTGCCCCGACATCCTTAATTATTACTTTTTTTTATTATTGTTGTTAT
ATGTACAATATAGGTTCATGGTCAGCATCGACAAGTCCGGCCCGTTCGCCTTCGCC
TTCTCGATATGGTGGTCATTTGTCTCGCAGTAAACGCACTCAACTGCCTTATCCCA
CATATGGGACAACCAGTCTATGTCAAAGATCACGATCGCCGAGTCCCGCTAGACTT
CAGGAGATGCGTGAACGAGACAGACTTGGTTttATGGGATTGATATGGGTACATCCT

TAGCACTTGTTATCTAAAGTCCAACAAGCTCAATCAAAAATCAACTTTAAACAATTTA
ACAAAAACAACTACTCAACTTGTTTTCTATATATGCTAGTATTTTCTTCTACACAAAT
TTCGACTTCAACCATCTCAATTTTAACAAACATCAAAGATCAGACATGCTTACTCGT
AAATCGTTAACTCGTAACTCGTATCTCGTAACTAACTTGTATTCGTACATAGTTGCC
AACCATGCTTAACATGTATACATGTATTGTCATGCCCAGACGTTCAATCAGAAAACA
AACAGTCAATCAATCAATCAATCAATCAATCAATCAATCAGCCAATCAGTCAACCAT
AAGCTTCAACAAATCGAATAGTTCAAATACTCTTTTCATACATCCAAACACAGCTCT
CAAAAATTCATTCAGATCTGCGATCTACCACACCGAAGAACTCATGATTATGTGACC
AACTATTCGTATTTCGTATTCGTACTCTTGTTACACTAGCAGCGTTTATGTTTTCATT
TGTCTAGAAGCCATCCTCGTAACATCCACCTATATTATAATCACATCTCGTATGTAT
GGTGCTGACATGTACAGACATATTCATCCTTGTCATTAAACTAACACACTGAATCTC
TGGCAAATGAACTGTTGAATAGCAATAGAGGAGTTTAAATATATACGATTTTTATGT
CAGGACTACTTTCGACGGACTCTTAGTAGAACACCAAGTACTGGTCACTTTTTAAAA
ACAGCATCGATGTGAGAGAAATGTGAATATATTATGGAACAGCAATGTGTAGGGGT
GAGATAAGTGAACATCATTTCCTAGTCTATTCCACAAAGAAGGTGAAGTTCGTAAG
CCATAGTATGTAGAACATATATATATATATATATAAATTTTTTTTTTTGTTCAAAATTGAC
CTCTGTCAGCCTGACCAGCACAGGATAAGACACTGACCAAGAATCTCTATTTAAAG
CTTGCTTCTTGTTCCAAAAAAAACACAAAAAAAAAAAACACCCAGTTACTACCACAGA
TATGCAGGCAATGTTCCAGTATTGTTTTTTGTACCAGTTTTGGGCGGTGAAATCGAC
TGAGCTACATACATACTCGTCTATGTAATATTATAACCAGAGGAATAATTTGGGTTG
CAGCCTTCGGGTGTTTCAGTATCTAGACGTGTGTGCATATTTATCTATCTTATCTAT
CTGTGAATGTGCGTCAATAGTATTCTGCACCATTTCTTAATCCATCCTTTTCGTTTTT
GCATAATTGATATCTCTATAGATCAAATCGAAATCAAAACAGTCACGCATGTTGCCA
TGAACAATTTAGCTGCCGACCGGTCATTACCCCAAATCATCCAACTACTTGTGTATA
TTTCTCTCAATACATTATATATAGCACTGTTGTTAATGTTTCATATTTTTCTAAGTTGT
ATAATTCTGTAGGGTGCTCAAATAATTCCACACGCTTATGAACGTAGATACAGTGAA
TGCATTTTTGGCATAGCAACAATTTTTAAACAACATTAAAGTTTAATATGTAAACTAC
GTTCTGATTGGGACATGTTCATCCTTTACACAATATCGTAAAATCACCATTAAATCG
TAATTGTGCAAAAAATATTAACTTCGAGTTTTGAAAGAACCTGTCAAATAAATCTTGT
TTCTATAAAGTTTGTATGTGTATTTTTAGAGATAACTAATGAAATCATAAATCGTTTA
ACGCTAACTTTGACCTGTACTTTCAGAATTCAATAGTCCTTACTGAGACTTAACAAA
ATCCCACTAAATGTCGCACAATAAAATTCAACATAAATCTGGTTACGATAAAGTCTG
TCTGATTGTCTTCAGTTATTAGATAGTTAGTCCTAATCTAATCCAAATGAACATGTCC
CATTCTATATTATGTTTCATATATTTGTATGTATGTTTTAAGTCCATGTTAATGTCAAT
GAAGAACGACTTAGGTACACTTTGTTCAGTATCATAATAGCAGAGTATTTTTGCAAC
GACTTATGCCCTAAACTGCACTTGTTCAGCACACAGGGAAGTAATTTTTCGATTTCC
CGCAGGTGTATTCGCATGTACAGCATAGTTACCCAACACTGGCCTCCCGAAGAGCC
GGAATCGGAAGACGCCTTCCTCCGACTCCCAGTAAACCGTCAACACTGCAGCTCA
AGCCAACCAATATCAATTTCCCGAAGCTCAATGCCAGTCCCACACATAGGAAGCGG
Agccaccaacttctccctgctgaagcaggccggcgacgtggaggagaaccccggccccatggtgagcaagggcgag
gagctgttcaccggggtggtgcccatcctggtcgagctggacggcgacgtaaacggccacaagttcagcgtgtccggcg
agggcgagggcgatgccacctacggcaagctgaccctgaagttcatctgcaccaccggcaagctgcccgtgccctggc
ccaccctcgtgaccaccctgacctacggcgtgcagtgcttcagccgctaccccgaccacatgaagcagcacgacttcttc
aagtccgccatgcccgaaggctacgtccaggagcgcaccatcttcttcaaggacgacggcaactacaagacccgcgcc
gaggtgaagttcgagggcgacaccctggtgaaccgcatcgagctgaagggcatcgacttcaaggaggacggcaacat
cctggggcacaagctggagtacaactacaacagccacaacgtctatatcatggccgacaagcagaagaacggcatca

aggtgaacttcaagatccgccacaacatcgaggacggcagcgtgcagctcgccgaccactaccagcagaacacccccc
atcggcgacggccccgtgctgctgcccgacaaccactacctgagcacccagtccgccctgagcaaagaccccaacga
gaagcgcgatcacatggtcctgctggagttcgtgaccgccgccgggatcactctcggcatggacgagctgtacaagtaa
AAGGAAGCGGAgccaccaacttctccctgctgaagcaggccggcgacgtggaggagaaccccggccccAtgg
cctcctccgaggacgtcatcaaggagttcatgcgcttcaaggtgcgcatggagggctccgtgaacggccacgagttcga
gatcgagggcgagggcgagggccgcccctacgagggcacccagaccgccaagctgaaggtgaccaagggcggcc
ccctgcccttcgcctgggacatcctgtcccctcagttccagtacggctccaaggcctacgtgaagcaccccgccgacatcc
ccgactacttgaagctgtccttccccgagggcttcaagtgggagcgcgtgatgaacttcgaggacggcggcgtggtgacc
gtgacccaggactcctccctgcaggacggcgagttcatctacaaggtgaagctgcgcggcaccaacttcccctccgacg
gccccgtaatgcagaagaagaccatgggctgggaggcctccaccgagcggatgtaccccgaggacggcgccctgaa
gggcgagatcaagatgaggctgaagctgaaggacggcggccactacgacgccgaggtcaagaccacctacatggcc
aagaagcccgtgcagctgcccggcgcctacaagaccgacatcaagctggacatcacctcccacaacgaggactacac
catcgtggaacagtacgagcgcgccgagggccgccactccaccggcgccTaa**ACCGGT**

**cac_34_GFP_mRFP1_remade**

**GCGGCCGC**caaaATGAGGCACAATGGCTCTCCACTGGCCAGATCTCCGAGTCCTCG
ACGGCGTGGCCATCAATACATACATCATGATATCGGGTTCTCCGATACCGTATCTAAT
GTTGTAGAGATGGTCAAGGAGACTCGTCATCCTAGGCATGGCAACAGTCATCCGCG
GTATCCAAGAGGTATTTATTGGCTCGACATCCATGGCATTTAATCATTAACTACTTTCA
ACTGACTATCGTGTGTGTGTGTGTGTGTGCTGGTTGTAAGCCAAATGATCTGGTTAG
GGATTCGTCAACATTATCCTGTTATCGAGTTGTCTGTCCTTGATGTTGCCCTTAATCA
CTTCGTATCCATATGAGATCGGCTTGCAAGGGGCACACACAGAAAACAAAGGGACT
TGGACATTCCAGGGATTTTTGATCTATTTTGTAGAACCGTAGTTATATACATATATACAT
ATATGTCTACAATTCCATAGCTATTAACGAAATCTTTTAATACAAATATATTATATTTCAAT
AATAATTTTATCGTTTCCGTTAAGTCGGTCCTCTATTGTCGTTTATCGTTTATCTCTTC
GAAAGCAATTTGAAAATCGCAAAAAACCAAAATGAAAAAAAAAAAAACAAAACAAAA
CAAAACAAAAGAAGAGAAGAGAAAATATGTTGCTAAGATCCAATACGTAAAATGTTGT
GCCCCGACATCCTTAATTATTACTTTTTTTTATTATTGTTGTTATATGTACAATATAGGTT
CATGGTCAGCATCGACAAGTCCGGCCCGTTCGCCTTCGCCTTCTCGATATGGTGGT
CATTTGTCTCGCAGTAAACGCACTCAACTGCCTTATCCCACATATGGGACAACCAGT
CTATGTCAAAGATCACGATCGCCGAGTCCCGCTAGACTTCAGGAGATGCGTGAACG
AGACAGACTTGGTTtATGGGATTGATATGGGTACATCCTTAGCACTTGTTATCTAAAG
TCCAACAAGCTCAATCAAAAATCAACTTTAAACAATTTAACAAAAACAACTACTCAAC
TTGTTTTCTATATATGCTAGTATTTTCTTCTACACAAATTTCGACTTCAACCATCTCAAT
TTTAACAAACATCAAAGATCAGACATGCTTACTCGTAAATCGTTAACTCGTAACTCGT
ATCTCGTAACTAACTTGTATTCGTACATAGTTGCCAACCATGCTTAACATGTATACATG
TATTGTCATGCCCAGACGTTCAATCAGAAAACAAACAGTCAATCAATCAATCAATCAA
TCAATCAATCAATCAGCCAATCAGTCAACCATAAGCTTCAACAAATCGAATAGTTCAA
ATACTCTTTTCATACATCCAAACACAGCTCTCAAAAATTCATTCAGATCTGCGATCTAC
CACACCGAAGAACTCATGATTATGTGACCAACTATTCGTATTTCGTATTCGTACTCTT
GTTACACTAGCAGCGTTTATGTTTTCATTTGTCTAGAAGCCATCCTCGTAACATCCAC
CTATATTATAATCACATCTCGTATGTATGGTGCTGACATGTACAGACATATTCATCCTT
GTCATTAAACTAACACACTGAATCTCTGGCAAATGAACTGTTGAATAGCAATAGAGG
AGTTTAAATATATACGATTTTTATGTCAGGACTACTTTCGACGGACTCTTAGTAGAACA

CCAAGTACTGGTCACTTTTTAAAAACAGCATCGATGTGAGAGAAATGTGAATATATTA
TGGAACAGCAATGTGTAGGGGTGAGATAAGTGAACATCATTTCCTAGTCTATTCCAC
AAAGAAGGTGAAGTTCGTAAGCCATAGTATGTAGAACATATATATATATATATAAATTTT
TTTTTTTGTTCAAAATTGACCTCTGTCAGCCTGACCAGCACAGGATAAGACACTGAC
CAAGAATCTCTATTTAAAGCTTGCTTCTTGTTCCAAAAAAAACACAAAAAAAAAAACA
CCCAGTTACTACCACAGATATGCAGGCAATGTTCCAGTATTGTTTTTTGTACCAGTTT
TGGGCGGTGAAATCGACTGAGCTACATACATACTCGTCTATGTAATATTATAACCAGA
GGAATAATTTGGGTTGCAGCCTTCGGGTGTTTCAGTATCTAGACGTGTGTGCATATT
TATCTATCTTATCTATCTGTGAATGTGCGTCAATAGTATTCTGCACCATTTCTTAATCCA
TCCTTTTCGTTTTTGCATAATTGATATCTCTATAGATCAAATCGAAATCAAAACAGTCA
CGCATGTTGCCATGAACAATTTAGCTGCCGACCGGTCATTACCCCAAATCATCCAAC
TACTTGTGTATATTTCTCTCAATACATTATATATAGCACTGTTGTTAATGTTTCATATTTT
TCTAAGTTGTATAATTCTGTAGGGTGCTCAAATAATTCCACACGCTTATGAACGTAGA
TACAGTGAATGCATTTTTGGCATAGCAACAATTTTTAAACAACATTAAAGTTTAATATG
TAAACTACGTTCTGATTGGGACATGTTCATCCTTTACACAATATCGTAAAATCACCATT
AAATCGTAATTGTGCAAAAAATATTAACTTCGAGTTTTGAAAGAACCTGTCAAATAAAT
CTTGTTTCTATAAAGTTTGTATGTGTATTTTTAGAGATAACTAATGAAATCATAAATCGT
TTAACGCTAACTTTGACCTGTACTTTCAGAATTCAATAGTCCTTACTGAGACTTAACA
AAATCCCACTAAATGTCGCACAATAAAATTCAACATAAATCTGGTTACGATAAAGTCT
GTCTGATTGTCTTCAGTTATTAGATAGTTAGTCCTAATCTAATCCAAATGAACATGTCC
CATTCTATATTATGTTTCATATATTTGTATGTATGTTTTAAGTCCATGTTAATGTCAATGA
AGAACGACTTAGGTACACTTTGTTCAGTATCATAATAGCAGAGTATTTTTGCAACGAC
TTATGCCCTAAACTGCACTTGTTCAGCACACAGGGAAGTAATTTTTCGATTTCCCGC
AGGTGTACCGCATGTACAGCATAGTTACCCAACACTGGCCTCCCGAAGAGCCGGAA
TCGGAAGACGCCTTCCTCCGACTCCCAGTAAACCGTCAACACTGCAGCTCAAGCC
AACCAATATCAATTTCCCGAAGCTCAATGCCAGTCCCACACATAGGAAGCGGAgccac
caacttctccctgctgaagcaggccggcgacgtggaggagaaccccggccccatggtgagcaagggcgaggagctgt
tcaccggggtggtgcccatcctggtcgagctggacggcgacgtaaacggccacaagttcagcgtgtccggcgagggcg
agggcgatgccacctacggcaagctgaccctgaagttcatctgcaccaccggcaagctgcccgtgccctggcccaccct
cgtgaccaccctgacctacggcgtgcagtgcttcagccgctaccccgaccacatgaagcagcacgacttcttcaagtccg
ccatgcccgaaggctacgtccaggagcgcaccatcttcttcaaggacgacggcaactacaagacccgcgccgaggtg
aagttcgagggcgacaccctggtgaaccgcatcgagctgaagggcatcgacttcaaggaggacggcaacatcctggg
gcacaagctggagtacaactacaacagccacaacgtctatatcatggccgacaagcagaagaacggcatcaaggtga
acttcaagatccgccacaacatcgaggacggcagcgtgcagctcgccgaccactaccagcagaacacccccatcggc
gacggccccgtgctgctgcccgacaaccactacctgagcacccagtccgccctgagcaaagaccccaacgagaagc
gcgatcacatggtcctgctggagttcgtgaccgccgccgggatcactctcggcatggacgagctgtacaagtaaAAGG
AAGCGGAgccaccaacttctccctgctgaagcaggccggcgacgtggaggagaaccccggccccAtggcctcctc
cgaggacgtcatcaaggagttcatgcgcttcaaggtgcgcatggagggctccgtgaacggccacgagttcgagatcgag
ggcgagggcgagggccgcccctacgagggcacccagaccgccaagctgaaggtgaccaagggcggccccctgccc
ttcgcctgggacatcctgtcccctcagttccagtacggctccaaggcctacgtgaagcaccccgccgacatccccgactac
ttgaagctgtccttccccgagggcttcaagtgggagcgcgtgatgaacttcgaggacggcggcgtggtgaccgtgaccca
ggactcctccctgcaggacggcgagttcatctacaaggtgaagctgcgcggcaccaacttcccctccgacggccccgta
atgcagaagaagaccatgggctgggaggcctccaccgagcggatgtaccccgaggacggcgccctgaagggcgag
atcaagatgaggctgaagctgaaggacggcggccactacgacgccgaggtcaagaccacctacatggccaagaagc
ccgtgcagctgcccggcgcctacaagaccgacatcaagctggacatcacctcccacaacgaggactacaccatcgtgg
aacagtacgagcgcgccgagggccgccactccaccggcgccTaa<u>**ACCGGT**</u>
